# BRD1 haploinsufficiency disrupts neurodevelopmental and metabolic homeostasis in a translational minipig model

**DOI:** 10.1101/2025.10.21.683618

**Authors:** Julie Grinderslev Donskov, Tue Fryland, Bolette Nicolaisen, Sanne Hage la Cour, Paula Rodrigo Martín, Steffi Pauwels, Lina Zühlsdorf, Dimitrios Pediotidis-Maniatis, Jacob Egemose Høgfeldt, Arne Mørk, Jane Hvarregaard Christensen, Ida Elisabeth Holm, Gregers Wegener, Simon Fristed Eskildsen, Torben Ellegaard Lund, Dora Gauballe, Jens Randel Nyengaard, Filip Ottosson, Madeleine Ernst, Aage Kristian Olsen Alstrup, Jannik Jakobsen, Anders Dupont Børglum, Per Qvist

## Abstract

Psychiatric disorders are complex conditions characterized by substantial overlap in genetic risk and shared biological mechanisms. However, how individual risk genes contribute to shared disease mechanisms and disorder-specific phenotypes remains poorly understood. BRD1 has emerged as a chromatin-associated regulator with transdiagnostic relevance across psychiatric disorders and a central role in gene regulatory networks enriched for psychiatric risk genes. To investigate the biological consequences of reduced BRD1 function in a translationally relevant system, we generated a minipig model harboring a monoallelic deletion in BRD1 and performed longitudinal neuroimaging together with behavioral and multi-omics profiling.

BRD1 haploinsufficient minipigs displayed normal growth and exploratory behavior but exhibited subtle age-dependent differences in motivational behavior. Despite the absence of overt developmental abnormalities, longitudinal neuroimaging revealed genotype-associated structural differences primarily involving the cerebral cortex and caudate nucleus, suggestive of altered neurodevelopmental trajectories. Integrated multi-omics analyses revealed striking convergence across transcriptomic and metabolomic datasets, identifying coordinated perturbations of mitochondrial function, redox regulation, and phospholipid metabolism across multiple brain regions. Notably, these molecular alterations were not restricted to the central nervous system, as peripheral multi-omics profiling revealed systemic metabolic alterations, including altered phospholipid composition and glucose metabolism.

Together, these findings indicate that BRD1 haploinsufficiency is associated with coordinated neurodevelopmental and metabolic alterations across brain and peripheral tissues. More broadly, this study provides systems-level insight into how a psychiatric risk gene influences interconnected neurodevelopmental and metabolic processes across multiple levels of biological organization and highlights the value of large-animal multi-omics models for translational neuropsychiatric research.

## Introduction

Psychiatric disorders are heritable and biologically complex conditions shaped by distributed genetic and environmental influences. Large-scale genomic studies have revealed substantial overlap in genetic risk across diagnostic categories, implicating shared molecular pathways involved in neurodevelopment, synaptic function, chromatin regulation, and cellular metabolism (1). These findings suggest that perturbations of fundamental biological processes contribute substantially to disease risk across traditional diagnostic boundaries. However, understanding how individual psychiatric risk genes influence biological systems across multiple levels of organization, from molecular regulation and brain development to systemic physiology, remains a major challenge.

The bromodomain-containing 1 gene (*BRD1*) has been repeatedly implicated in psychiatric disorders (2–5), with large-scale genome-wide association studies (GWAS) demonstrating cross-diagnostic associations (6,7), including genome-wide significant association observed for schizophrenia (SZ) (8). Rare variant analyses have further identified disruptive mutations in SZ and autism spectrum disorder (ASD) patients (5,9–13). *BAD1* encodes a chromatin-associated scaffold protein that forms part of histone acetyltransferase complexes (14,15) and influences the expression of large gene sets (15,16)Click or tap here to enter text.. (15)Importantly, both the chromatin and protein interactome of BRD1 is enriched with SZ and bipolar disorder (BPD) risk genes (16), supporting its role in molecular regulatory systems relevant to psychiatric disease biology.

*BAD1* is abundantly expressed in the developing brain across species (4,20,21) and plays important roles in early neurodevelopment, synaptic function, and neuronal network formation(22–24). In accordance, experimental rodent models with targeted disruption of *Brd1* display behavioral phenotypes relevant to psychiatric disorders, including altered affective behavior, cognitive dysfunction, selective sensitivity to psychostimulants, and behaviors indicative of hyperarousal (25–28). (25,28) These phenotypes are accompanied by gender-specific regional alterations in associated neurotransmitter systems (25,28) and structural brain abnormalities have been detected in male *Brd1*^+/−^ mice, including reduced cortical, striatal and amygdaloid volumes (29). At the molecular level, whole transcriptome profiling of selected brain regions in *Brd1*^+/−^ mice has revealed dysregulation of neurodevelopmental and signal transduction pathways enriched for psychiatric risk genes (25), while recent experimental work further suggests that BRD1 deficiency impairs mitochondrial bioenergetics (30) Collectively, these findings support a broader role for BRD1 in coordinating neurodevelopmental and metabolic processes relevant to psychiatric vulnerability.

Although rodent models have provided valuable insights into psychiatric disease mechanisms, important translational limitations remain, particularly with respect to neurodevelopmental trajectories, cortical organization, and systems physiology (31). The Göttingen minipig represent an attractive non-primate translational model (32) owing to its (32) large, gyrencephalic brain (33), human-like neuroanatomy (34), and prolonged postnatal maturation (35), which more closely parallels human neurodevelopment (36,37). In addition, pigs display physiological and endocrine characteristics that resemble those of humans, including aspects of stress responsivity and metabolic regulation (38).

Here, we report the generation and characterization of a genetically modified pig model with a targeted disruption of *BAD1*. Using longitudinal neuroimaging together with transcriptomic, metabolomic, lipidomic, and behavioral profiling, we investigated the impact of *BAD1* hapoinsuficiency across brain and peripheral tissues. Through this systems-level approach, the study provides insight into how disruption of a psychiatric risk-associated chromatin regulator influences neurodevelopmental, metabolic, and neurological processes in a translationally relevant large-animal model.

## Materials and Methods

### Animals

All experiments were approved by the Danish Animal Experiments Inspectorate (license no. 2019-15-0201-01633/CHNER) in accordance with 2010/63/EU and conducted at Aarhus University and Aarhus University Hospital. The study was reported in adherence to ARRIVE guidelines.

#### Animal housing and husbandry conditions

Göttingen Minipigs (Ellegaard Göttingen Minipigs A/S, Denmark) (39) were housed and cared for in accordance with the ministerial orders of the Danish Animal Experiments Inspectorate on genetically modified animals and the EU Directive 2010/63/EU for animal experiments. Göttingen Minipigs were fed a restricted pellet diet (SDS SMP (E) SQC Minipig diet; Special Diet Services, UK, purchased from Scanbur, Denmark) with tap water available ad libitum. The pigs were housed in pairs at 20–22 °C with a relative humidity of 50–55%, and 12hr light/dark day–night cycle. A climate system ensured fresh air was supplied eight times an hour. Animals were introduced to test facilities and acclimatized prior to experiments.

#### Anesthesia and animal preparation for imaging

The overnight fasted minipigs were transported to the scanning facility at Aarhus University Hospital and here treated with approximately 1.3 mg/kg midazolam and 6.3 mg/kg s-ketamine IM. After some minutes of sedation, an ear vein catheter was placed, and anesthesia was induced with approximately 1.3 mg/kg midazolam and 3.1 mg/kg s-ketamine IV as a bolus. After tracheal intubation, the anesthesia was maintained with 1½–3% sevoflurane (dosing after effect). The pigs were mechanically ventilated 15 times per minute with approximately 8-9 mL/kg of an oxygen-medical air (1: 2.2) mixture (40). Pulse, SatO2 and body temperature were measured, as well as interdigital reflexes controlled during the anesthesia. The pigs were placed in the scanner as described by others (41).

### *BRD1* transcript sequences and isoforms in cortical tissue from the Göttingen Ellegaard minipig

Guided by orthologue genomic *BRD1* sequence data from human, (Homo sapiens), chimpanzee (Pan troglodytes), mouse (Mus musculus), dog (Canis lupus familiaris), sheep (Ovis aries) and swine (Sus scrofa), PCR primers were designed targeting swine *BRD1* expressed sequence tags (EST) conserved across species, covering the length of the 12 human *BRD1* exons. For the PCR reactions the “Q5® Hot Start High-Fidelity DNA Polymerase”-kit was used following manufacturer’s instruction. Primers and settings for the thermocycler can be seen in **Table S1 and 2**. PCR products were separated by length using electrophoresis and fragments were analyzed using Sanger sequencing.

### Single guide RNA design and vector construction

Single guide RNA (sgRNA) was designed using the ZiFit (42) CRISPR design tool targeting exon 5 and 6 of the porcine *BRD1* gene. Exons 5 and 6 were chosen because of their relatively long sequences and similarity to the human exons 5 and 6. The exon 5 target sequence was: GATGCACCTGCCCGAGCGGC, and for exon 6: GGAG-TCAGGTACTGGCGGCC, and the following primers were used in pairs: 5’ CAC CGA TGC ACC TGC CCG AGC GGC 3’ – 5’ AAA CGC CGC TCG GGC AGG TGC ATC 3’ and 5’ CAC CGG AGT CAG GTA CTG GCG GCC 3’ – 5’ AAA CGG CCG CCA GTA CCT GAC TCC 3’. sgRNA was introduced into CRISPR-Cas9 transposon vectors (pX330-U6-Chimeric_BB-CBh-hSpCas9 plasmid, Addgene no. 42230) using a slightly modified version of the Zhang Lab General Cloning Protocol (43). Followed by transformation and selection of Ampicillin resistant cells, subcultured colonies were purified with NucleoBond Xtra plasmid purification kit from Macherey-Nagel.

### Transfection and selection of cell clones

Fibroblasts were cultured from neonate ear biopsies derived from a neonate Göttingen master minipig (498) containing two integrated heterogeneous loxP sites. The cells were grown in Dulbecco’s Modified Eagle Medium (DMEM) with 15% fetal calf serum (Sigma-Aldrich, batch F7524; 042M3396), 1% penicillin/streptomycin (P/S), and 1% glutamine (Sigma-Aldrich, St. Louis, MO, USA) to 50% confluence and passaged for further expansion prior to freezing of aliquots. Fibroblasts were seeded at 30% confluence and transfected using 700 ng Blue-pyro plasmid (**Figure S1**), 1200 ng pX330-U6-Chimeric_BB-CBh-hSpCas9_BRD1_5/6 plasmid and 100 ng plasmid encoding the CRE recombinase using 6 μL TurboFect lipofectamine (Thermo Fisher Scientific, Waltham, MA, USA). Following puromycin selection of plated cells, DNA was purified with the Maxwell DNA purification kit (Promega). Mutations at the target site were analyzed using Sanger sequencing of PCR products and the ICE v2 CRISPR Analysis Tool.

### Handmade cloning, embryo culture, and transfer

Three cell colonies with validated disruptive monoallelic *BRD1* mutations were chosen for somatic cell nuclear transfer (SCNT) to create Göttingen Minipigs with monoallelic disruption to *BRD1*. Selected colonies were pooled and grown for nine days prior to SCNT by handmade cloning (HMC). HMC was performed as described (44). Reconstructed embryos were incubated and cultivated for 5–6 days before surgically transferred into Danish landrace surrogate sows. Only one surrogate sow delivered piglets. They were delivered by natural farrowing on day 115 resulting in four viable male F0 generation genetically modified *BAD1*^+/−^ minipigs raised by their surrogate mother. Mating between F0 *BAD1*^+/−^ boars and WT Göttingen Minipigs subsequently produced two litters of F1 minipigs, specifically five males (five *BAD1*^+/−^ and zero WT piglets) and seven females (four *BAD1*^+/−^ and three WT piglets). Since no WT male piglets were born in the F1 generation, this study utilized only female *BAD1*^+/−^ and WT minipigs.

### Behavioral assessment of minipigs

Three WT and four *BAD1*^+/−^ female minipigs were subjected to a series of behavioral trials.

#### Open field

The open field (OF) test was used to measure exploratory behavior and activity level under different circumstances. Trials were carried out in a 200*270 cm arena with four walls and one entrance. The arena floor was divided with duct tape into twelve squares of equal size, with a letter representing each square, as a visual aid for tracking and scoring their movements. Video recordings with a duration of 9-10 minutes were acquired from four female *BAD1*^+/−^ minipigs and three female WT minipigs. On day 1, the pigs were introduced to the arena for the first time and their response to the new surroundings was assessed (OF novelty). On day 3, the pigs were presented to a foreign object (ball) in the arena (OF novelty + object). Here we assessed their reaction to a foreign object in a familiar setting. On day 6, no object was present in the arena, and their baseline level of activity in familiar surroundings was assessed (OF baseline). The time spent in any given square and number of crossings between squares were scored for all subjects. “Crossing” was observed when both shoulders of a pig entered a different square. Scoring began the moment the experimenter had left the arena, and the pig was safely inside the arena with the door closed. All video recordings were analyzed manually by an observer blinded to genotype and minipig ID using the Noldus Ethovision video tracking software.

#### Sucrose preference

Female *BAD1^+/−^* and WT minipigs were tested in a sucrose preference test. All pigs were single-housed for a minimum of one hour prior to testing proceeding into testing. For a period of two hours, their regular water supply was halted, and two “non-spill” water troughs containing either one liter of sucrose water or plain water were placed in each pig enclosure. When the test was over, troughs were removed, pigs regrouped, and the remaining amount of water in each trough was measured. Pigs were single-housed and familiarized with the new water troughs a few days prior to testing, and their “regular” water intake estimated.

### Magnetic resonance imaging

The minipigs were scanned on a Magnetom Prisma 3T MRI system (Siemens Healthcare, Erlangen, Germany) using a 32-channel head coil. Whole head 3D T1-weighted MP2RAGE (magnetization-prepared two rapid gradient echo acquisitions) (Marques et al., 2010) images were acquired (600 µm isotropic voxels without gap, TR/TE=5000/2.72 ms, TI_1_=700 ms, TI_2_=2500 ms, flip-angle_1_=4°, flip-angle_2_=5°, FOV=234×250 mm^2^ and 125 mm in the sagittal direction). To increase signal-to-noise ratio, two acquisitions were carried out back-to-back and averaged in a postprocessing step. Images were denoised (45) bias field corrected (46) and cropped to the brain. For each image, a brain mask was created using intensity classifications followed by morphological operations and manual corrections if necessary. Age-specific templates were created for each time point (6 months, 9 months and 14 months) using an iterative framework nonlinearly aligning the images following the procedure described in Fonov et al. (47). As reference space we used a template based on in vivo MRI of the neonatal piglet available from the Pig Imaging Group at University of Illinois ((https://pigmri.illinois.edu)) (48). Atlas labels from the Illinois template were transformed to the age-specific templates followed by manual correction and manually adding segmentation of amygdala. Atlas labels were then warped from the age-specific templates to the corresponding native images using the transformation matrices and deformation fields generated during template creation. Whole brain and ROI volumes were calculated in native space.

### Biosample collection and blood glucose measure

Peripheral blood samples were collected from the jugular venipuncture at 6, 9, and 14 months of age. A total of 18 ml was drawn per time point, divided equally into two 9 ml tubes for plasma and serum preparation. For plasma, tubes were gently inverted several times, kept on ice, and centrifuged at 1500 × g for 15 min at 4 °C. The supernatant plasma layer was carefully transferred to cryovials and stored at −80 °C until analysis. For serum, blood was allowed to clot for 30 min at room temperature, then centrifuged under identical conditions, and the resulting supernatant was aliquoted and stored at −80 °C.

All pigs were fasted for a minimum of 16 hours prior to sampling to reduce variability in metabolic measures. Blood glucose was measured at age 14 months using an Accu-Chek Instant Blood Glucose Meter clinical glucose meter. At 14 months, pigs were euthanized and biosample collection was performed immediately postmortem. Whole brains were extracted and weighed. The right cerebral hemisphere was preserved by immersion fixation in formalin, while the left hemisphere was weighed and used for regional dissections. The cerebrum was separated from the mesencephalon, and the brain-stem and mesencephalon were snap-frozen. The cerebellum was isolated, sliced into ∼1 cm thick blocks, and regions of interest were dissected using a standardized coronal slicing template placed on a chilled plate. From the left hemisphere, the following brain regions were collected by free-hand dissection and snap-frozen: anterior cingulate cortex (aCC), lateral prefrontal cortex (LPFC), caudate nucleus (NC), putamen (Pu), nucleus accumbens (NAc), hypothalamus, amygdala (AMG), and hippocampal subfields (CA1, CA3, dentate gyrus). Tissue was also collected from the cerebellar vermis (spinocerebellum).

### Quantification of neurotransmitters by HPLC

After weighing, aCC, AMG, NC and NAc samples were homogenized in ice-cold 0.05M HClO4 (Sigma-Aldrich), and centrifuged (20000×g) for 30 min. at 4°C. Supernatants were filtered (0.22 µm column; Millipore, Billerica, USA), and were separated by HPLC (ODS 150×2 mm column; flow rate 0.2 ml/minute). Dopamine and serotonin were electrochemically detected (E2=200 mV) by Coulochem-III (Thermo scientific, Sunnyvale, USA).

### Transcriptomic profiling using RNA sequencing

RNA was extracted using Maxwell-16 instrument system and LEV simplyRNA Tissue Kit (cat. no. AS1280, Promega, Madison, USA) according to manufacturer’s protocol, except tissue homogenization was performed with a handheld OMNI tissue homogenizer that was rinsed in nuclease-free water and ethanol between each sample. Furthermore, 10 μL of DNAse 1 were added consequently to the 4th well instead of the 5 μL stated in the protocol. Agilent 2100 Bioanalyzer (Agilent technologies, SantaClara, USA) confirmed the quality of RNA with a mean RNA Integrity Number (RIN) of 7.87 (SD 0.26). For all samples, libraries were prepared using Beijing Genomics Institute (BGI) library preparation kits and protocols and sequencing performed on the BGISEQ-500 platform. A minimum of 10 million clean 50 bp single-end reads were generated for each sample. Reads that passed quality control (more than 90% bases having less than 1% sequencing error; No ambiguous bases) were aligned to pig genome (Sus scrofa) by HISAT2 (version 2.1.0) (49) and counted by StringTie (version 1.3.4) (50). Differentially expressed genes (DEGs) were identified by DESeq2 and reported after Benjamini-Hochberg false discovery rate (FDR) (5% correction) (14, 16) or as nominally significant DEGs (p < 0.01). Gene-set enrichment analysis and overrepresentation analysis were done using ClusterProfiler(v4.12.6) (51).

### Metabolomic and lipidomic profiling

Metabolite profiling was carried out at the Section for Clinical Mass Spectrometry, Danish Center for Neonatal Screening, Department of Congenital Disorders, Statens Serum Institut, Denmark. A total of 28 brain tissue samples and 20 plasma samples were submitted to untargeted LC-MS/MS based metabolomics profiling as described below. Lipid analysis on 21 serum samples was performed by cmbio (https://cmbio.io/).

#### Sample preparation

Brain sections exclusively from the left hemisphere were prepared for MS-analysis using a standard tissue homogenization protocol: 30 mg of brain tissue was homogenized in ice-cold 400 μg 0.1% formic acid in MeOH using a Precellys 24 instrument (2×15 s at 5000 rpm, two steel beads per sample), followed by centrifugation (10 min 10 000 ×g 4 °C). 300 μL of the supernatant was transferred to a new tube and dried out for 2 hours at 50 °C in a vacuum centrifuge. Samples were extracted in 100 µL of 5% solvent B (49.9% methanol, 49.9% acetonitrile and 0.2% formic acid) and 95% solvent A (99.8% water and 0.2% formic acid), shaken during 30 minutes at 650 rpm at room temperature and 75 µL were transferred to a 96-well plate. Pools of all brain tissue samples were analyzed for quality control purposes.

Plasma samples were vortexed and 100 µL of each sample was distributed on a 96 deep well plate. 300 µL of cold methanol/acetonitrile (1:1) was then added and samples were shaken for 15 minutes at 900 rpm at room temperature. Samples were then frozen at −20°C for 1 hour and centrifuged for 30 minutes at 4°C. 15 µL were then transferred to a new 96-well plate and dried under nitrogen for 1 hour at 60 L/min. Samples were then reconstituted in 100 µL 5% solvent B and 95% solvent A and shaken for 15 min. at 600 rpm at room temperature. Finally, the plate was centrifuged for 10 min. at 3000 rpm and 4°C. Pools of all plasma samples were analyzed for quality control purposes.

#### Mass spectrometry analysis (Metabolomics)

Mass spectrometry analysis was performed using a timsTOF Pro mass spectrometer coupled to a UHPLC Elute LC system, Bruker Daltonics (Billerica, MA, US). The analytical separation was performed on an Acquity HSS T3 (100 Å, 2.1 mm × 100 mm, 1.8 µm) column (Waters, Milford, MA, US). The analysis started with 99% solvent A for 1.5 min, thereafter a linear gradient to 95% solvent B during 8.5 min followed by an isocratic condition at 95% mobile phase B for 2.5 min before going back to 99% mobile phase A and equilibration for 2.4 min. Metabolomics preprocessing was done using the Ion Identity Molecular Networking workflow in MZmine (52) (version 3.3.9). Before statistical analysis, metabolite features present in less than 20% of the samples were removed and missing values were replaced by a random number between zero and the lowest detected value for each feature. For the plasma metabolomics, this resulted in a final dataset with a total of 1221 metabolite features measured, while the corresponding numbers for the brain sample metabolomics was 2530 metabolites. Subsequently all metabolites underwent mean centering and univariance scaling. Annotation of metabolite features was performed using mass spectral molecular networking through the GNPS Platform (53,54), and *in silico* annotation using Sirius+CSI:FingerID (55) and deep neural networks in CANOPUS (56).

#### Mass spectrometry analysis (Lipidomics)

The analysis was carried out using a Thermo Scientific Vanquish LC coupled to Thermo Q Exactive HF MS. Ionization was performed in positive and negative ionization mode using an electrospray ionization interface. The chromatographic separation of lipids was carried out on a Waters® ACQUITY Charged Surface Hybrid (CSH™) C18 column (2.1 × 100 mm, 1.7 μm). The column was thermostated at 55°C. The mobile phases consisted of (A) Acetonitrile/water (60:40) and (B) Isopropanol/acetonitrile (90:10), both with 10 mM ammonium formate and 0.1% formic acid. Lipids were eluted in a two-step gradient by increasing B in A from 40 to 99% over 18 min. Flow rate was 0.4 ml/min. Peak areas were extracted using Compound Discoverer 3.3 (Thermo Scientific). Identification of compounds were performed at four levels. For this study, only Level 1 annotations were retained, corresponding to lipids identified by matching accurate mass (within 3 ppm), retention time against in-house authentic standards, and MS/MS spectral confirmation. Lipid annotations were matched to LIPID MAPS.

#### Data analysis

Preprocessed metabolomics (brain and plasma) and lipidomics (serum) datasets were analyzed in MetaboAnalyst 5.0 (metaboanalyst.ca). For brain-derived data, genotype-dependent differences between *BAD1*^⁺/⁻^ and WT pigs were evaluated by univariate analysis (t-tests, p < 0.05). Pathway enrichment analyses were performed using the *Enrichment Analysis* module of the MetaboAnalyst platform (v5.0) (1). All annotated compounds were uploaded as the reference metabolome. Nominally altered metabolites (p < 0.05) were then used as the query list. Over-representation analysis was conducted against the Small Molecule Pathway Database (SMPDB), and pathway significance was assessed using the hypergeometric test with Holm adjustment for multiple testing. Joint pathway analysis was then performed using the *Pathway Analysis - Joint Pathway Analysis* module from the MetaboAnalyst platform (v6.0). Nominally altered metabolites and differentially expressed genes per tissue (p < 0.05), together with their respective log fold-change (logFC) scores, were included in the analysis. Pathway enrichment was conducted against the integrated metabolic pathways database, which includes pathways containing both metabolites and metabolic genes. Enrichment analysis was performed using the hypergeometric test, pathway topology analysis was based on degree centrality and integration method was set to combined queries. For blood metabolomics and lipidomics data, a linear model was applied with genotype as the main factor, age as a covariate, and pig ID as a blocking factor to account for repeated measures. Significance was evaluated using both raw and FDR-adjusted p-values.

## Results

### Generation of a *BRD1* haploinsufficient minipig model

To identify suitable target sequences for genomic editing of the previously uncharacterized Göttingen minipig *BAD1* gene, we first sequenced *BAD1* transcripts in fibroblasts from a Göttingen minipig (**Figure S2A**). Multiple transcript variants were identified, including a major isoform closely resembling human *BAD1* exon organization (**Figure S2B**). Despite differences in exon composition, all detected porcine *BRD1* transcripts retained the conserved bromodomain, PWWP domain, and PHD-finger domains present in human *BAD1*, supporting functional conservation. To target all transcripts, sgRNAs were designed against a sequence shared across all detected transcript variants.

Following CRISPR-Cas9 mediated genomic editing in fibroblasts isolated from a Göttingen minipig, a clone carrying a 14bp monoallelic deletion in exon 6 of *BAD1* was selected for hand-made cloning (**Figure A**). The deletion introduces a premature stop codon predicted to promote degradation of the altered transcript through non-sense mediated decay.

Gene-edited fibroblast nuclei were subsequently transferred into enucleated oocytes derived from a landrace pig using somatic cell nuclear transfer (SCNT) (**Figure 1B**). Supporting the efficiency of the selected genomic editing strategy, fibroblasts from first generation *BAD1*^+/−^ minipigs (F0) displayed significantly reduced BRD1 protein levels compared to wild-type (WT) controls (**Figure S3A-E**, BRD1-S: F = 9.39, *p* = 0.014, BRD1-L: F = 9.32, *p* = 0.014). Subsequent breeding with WT female Göttingen Minipigs generated mixed litters of WT and *BAD1*^+/−^ offspring.

**Figure 1.**
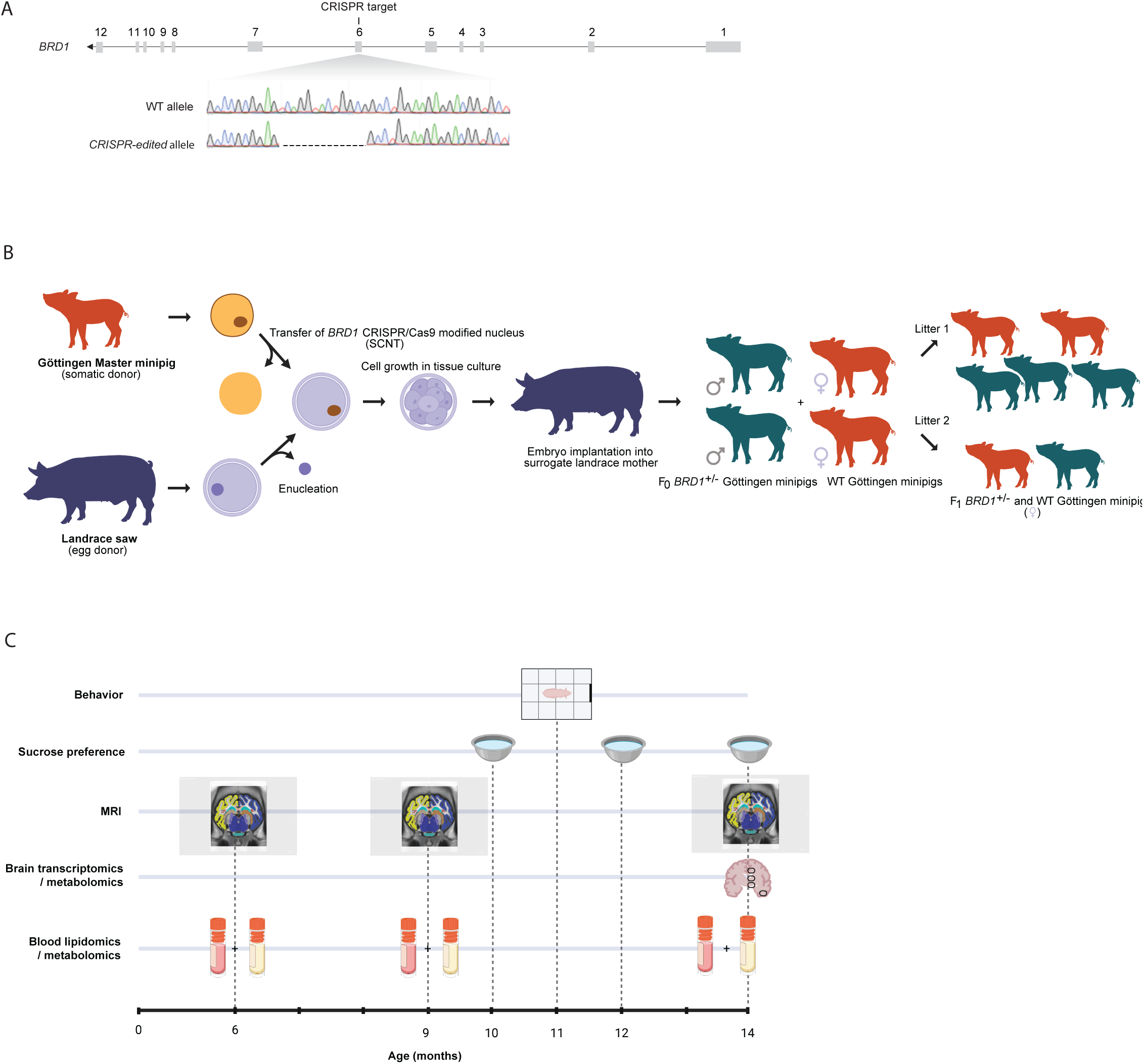
Generation of *BRD1^+/−^* minipig model and study design. **A.** Validation of monoallelic *BAD1* deletion in minipig fibroblasts by Sanger sequencing. Targeting all identified porcine *BAD1* transcripts, sgRNAs for genomic precision editing were designed for the sequence downstream of exon 4. CRISPR-Cas9-mediated genomic editing resulted in a clone carrying a 14bp monoallelic deletion in exon 6 of the porcine *BAD1* gene, which was selected for hand-made cloning. **B.** Hand-made cloning of Göttingen Minipigs. CRISPR-modified nuclei from fibroblasts were isolated and transferred to an enucleated oocyte derived from a landrace pig by SCNT. Embryo implantation into a surrogate landrace pig produced the first generation (F0) of *BAD1^+/−^* minipigs, while mating male F0 pigs with female WT pigs produced two litters of second-generation (F1) *BAD1^+/−^* minipigs. Of these, three WT and four *BAD1^+/−^* female pigs served as experimental subjects for this study. **C.** Study design showing timepoints for collected data or samples for experiments on female minipigs. Whole and regional brain MRI was performed at 6, 9 and 14 months of age; metabolomics (plasma and tissue) and lipidomics (serum and tissue) at 6, 9 and 14 months; sucrose preference at 10, 12 and 14 months; open field and object recognition at 11 months; bulk RNAseq of tissues from four brain regions at 14 months (immediately following euthanization).

To assess the neurodevelopmental and molecular impact of *BAD1* haploinsufficiency, we initiated longitudinal sampling of biofluids, repeated brain imaging, and terminal tissue collection in the first filial generation (F1) littermates (**Figure 1C**). Because no WT males were born in the F1 generation, all analyses were restricted to female pigs to avoid sex-related confounding.

### *BRD1* haploinsufficient pigs exhibit normal growth and exploratory behavior but subtle alterations in motivational response

No gross abnormalities were observed in the F1 litter, and *BAD1*^+/−^ pigs displayed normal growth trajectories throughout development (**Figure S4**). To assess behavior, pigs were subjected to repeated open field (OF) testing, including exposure to a novel object. Both *BAD1*^+/−^ and WT pigs engaged readily in exploratory behavior across sessions, although substantial inter-individual variability was observed (**Figure S5A-B**). No consistent genotype-dependent differences in general locomotor or explorative activity were detected during initial OF exposure (**Figure S6A-B**), novel object testing (**Figure S6C-D** and **Figure S7A-B**), or repeated arena exposure (**Figure S6E-F** and **Figure S8A-B**).

Given the variability observed in arena-based behavioral measures, we additionally assessed motivational behavior using a sucrose preference test (57). While overall sucrose preference was similar in younger ages, *BAD1*^+/−^ pigs displayed a modest age-dependent reduction in sucrose preference that became apparent from 10 months of age (**Figure 2**, F = 10.42, *p* = 0.023).

**Figure 2.**
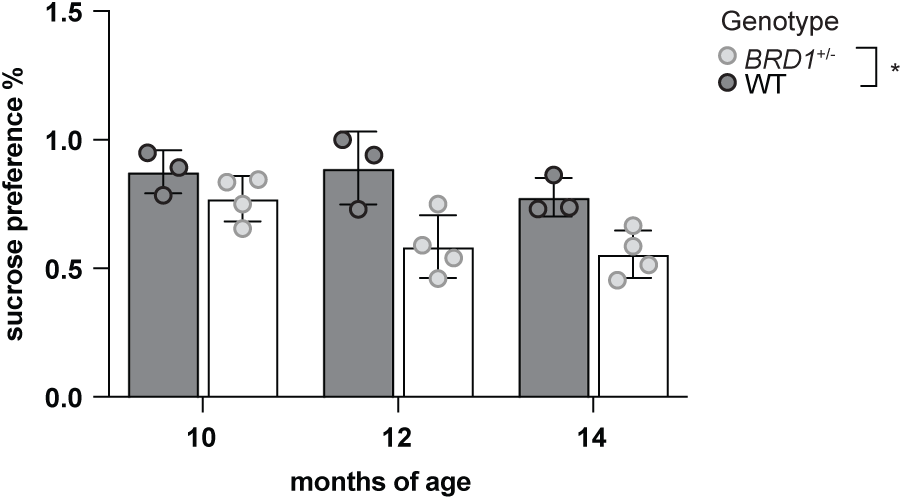
Sucrose preference measured at 10, 12 and 14 months of age. Bars represent genotype mean±SD.

### Longitudinal brain imaging uncovers *BRD1* haploinsufficiency-associated structural changes

To investigate the impact of *BAD1* haploinsufficiency on brain development, we performed longitudinal structural brain imaging on female F1 littermates at 6, 9, and 14 months of age. Initial inspection of raw volumetric data indicated a substantial contribution of litter identity to overall variance in brain volume estimates (**Figure S9**). Hence, to reduce litter-associated confounding effects, batch correction was applied prior to downstream analyses.

Following correction, principal component analysis (PCA) of regional brain volumes revealed geno-type-associated clustering of *BAD1^+/−^* and WT pigs along the PC2 axis, supporting genotype-associated differences in brain structural organization (**Figure 3A**). While *BAD1*^+/–^ pigs showed a trend toward smaller total brain volume compared to WT controls, considerable inter-subject variability was observed and the difference did not reach statistical significance (**Figure 3B**). However, to account for variation in overall brain size, regional volume estimates were normalized to total brain volume for subsequent analyses.

**Figure 3.**
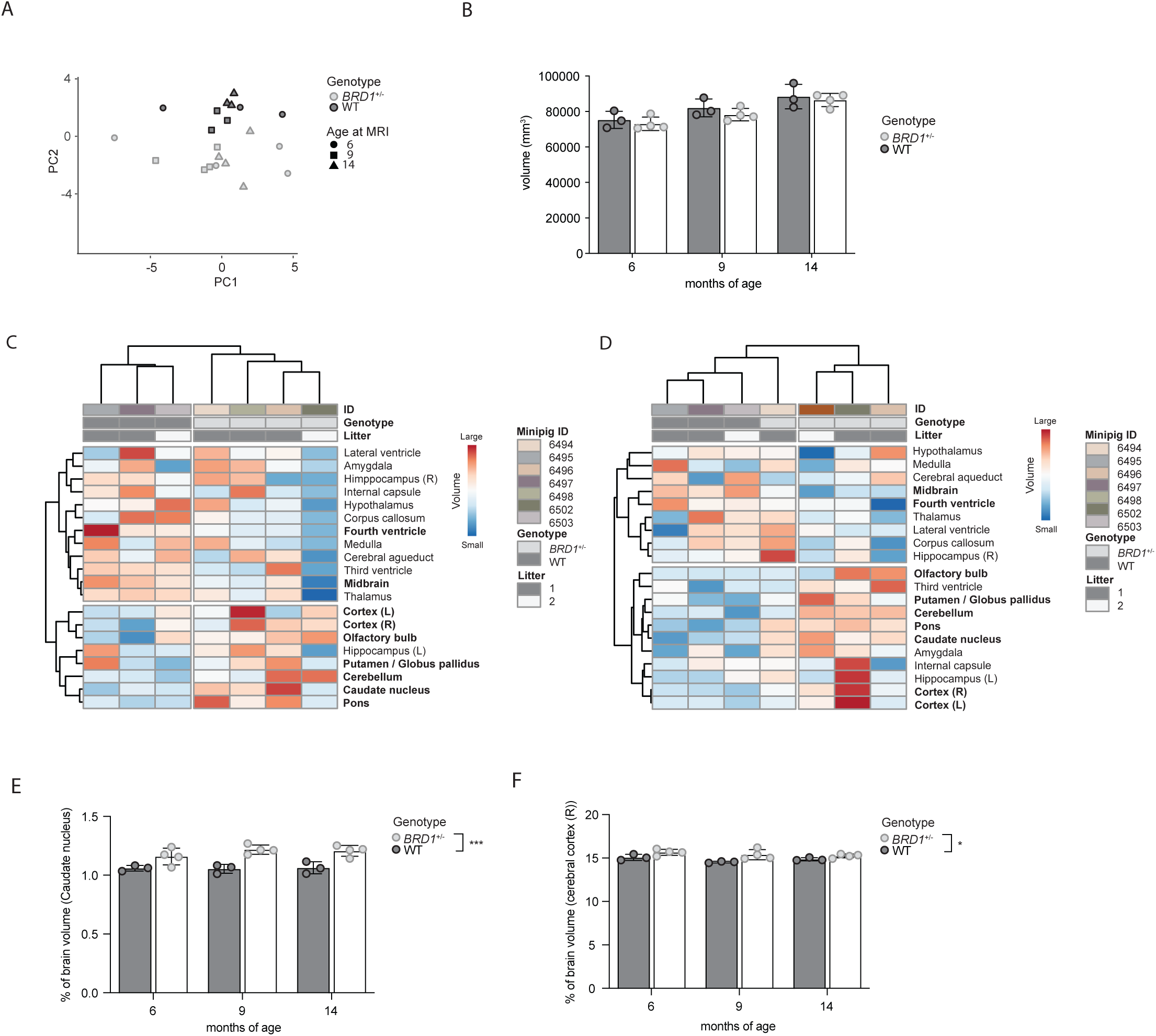
Structural brain imaging of *BRD1^+/−^* and WT minipigs. Minipigs underwent MRI at 6, 9 and 14 months of age and volumes of whole brain and 20 brain regions were assessed for each timepoint by manual segmentation. **A.** Principal component analysis based on all regional brain volumes. **B.** Minipig whole brain volume. **C–D.** Heatmaps showing hierarchical clustering of normalized and litter-corrected regional brain volumes at 6 months **(C)** and 9 months **(D)** of age, showing consistent regional changes in certain brain regions over time. **E.** Development of caudate nucleus volumes over time. Bars represent the mean proportion of total brain ± SD. **F.** Development of right cerebral cortex volumes over time. Data represent the mean proportion of total brain volume ± SD.

Region-wise analyses revealed distributed region-specific volumetric remodeling across developmental time points in *BAD1*^+/−^ pigs. Hierarchical clustering and heatmap visualization demonstrated consistent genotype-associated volumetric changes involving multiple brain regions, including the cerebral cortex, caudate nucleus (NC), olfactory bulb, pons, and cerebellum (**Figure 3C-D** and **Figure S10**). Several regions, including the cerebral cortex and NC, displayed increased relative volumes in *BAD1*^+/−^ pigs, whereas reductions were observed in the midbrain and fourth ventricle. Notably, the magnitude of regional effects varied across developmental stages. Statistical testing identified significant genotype effects in the NC (**Figure 3E**, F = 52.85, *p* = 0.0008), and right cerebral cortex (**Figure 3F**, F = 8.41, *p* = 0.03).

### *BRD1* haploinsufficiency drives transcriptomic changes and metabolic dysregulation in the brain

To investigate the molecular impact of *BAD1* haploinsufficiency across brain regions, we performed transcriptomic profiling in postmortem samples from NC, nucleus accumbens (NAc), amygdala (AMG), and the anterior cingulate cortex (aCC). Bulk RNA sequencing revealed substantial region-specific transcriptomic alterations, with the largest changes observed in AMG.

In total, 88 genes were differentially expressed (DEGs; padj.<0.05) in the AMG of *BAD1*^+/−^ pigs compared to WT (**Figure 4A** and **Table S3**). These included genes involved in excitatory and inhibitory neurotransmission and synaptic signaling, several of which have previously been implicated in psychiatric disorders, including GRM3 (58), *GAM4* and *GABAE* (59). More limited transcriptomic changes were observed in the aCC, NAc, and NC, where eight, six, and five DEGs were identified, respectively (**Figure 4B-D** and **Table S4-S6).** Across regions, pathway enrichment analyses converged on biological processes related to synaptic signaling, cellular organization, mitochondrial function, RNA and protein metabolism, and broader neuro-metabolic regulatory systems (60–62) (**Figure 4E-I, Figure S11-S13**). Notably, pathways linked to oxidative phosphorylation, selenoamino acid metabolism, translation, RNA splicing, and neural system function were repeatedly enriched across multiple regions (**Figure 4I** and **Figure S11-S13**).

**Figure 4.**
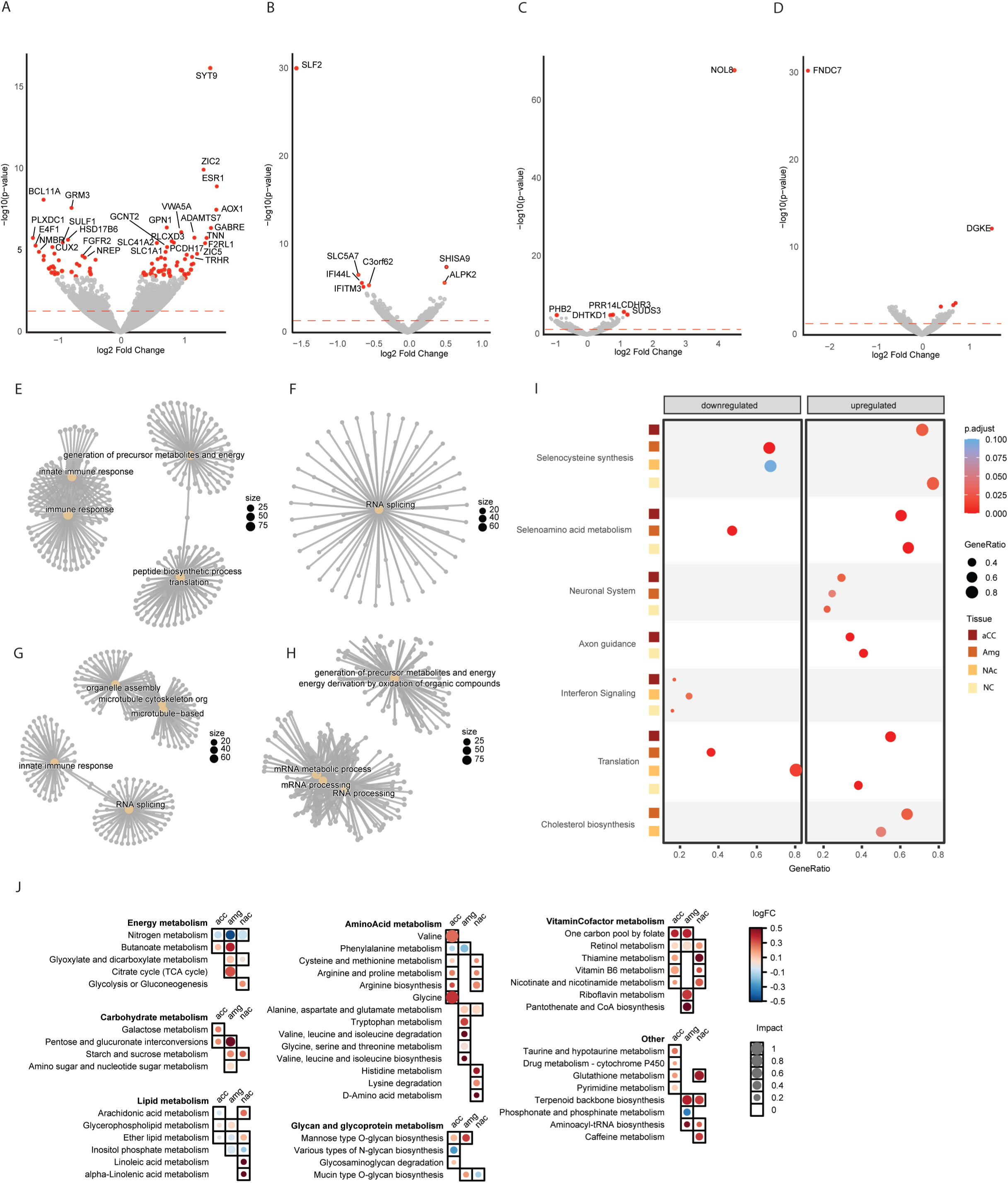
BRD1 haploinsufficiency drives transcriptomic changes and disrupts metabolic homeostasis in the brain. A–D. Volcano plots presenting differentially expressed genes in amygdala **(A),** anterior cingulate cortex **(B),** nucleus accumbens **(C)** and caudate nucleus **(D)** of *BAD1^+/−^*vs. WT minipigs. The –log₁₀(adjusted p-value) is plotted against the log₂(fold change) for each gene. Genes meeting the thresholds for FDR-corrected statistical significance (adjusted p-value < 0.05) are highlighted in red and non-significant genes in gray. **E-H.** Gene network visualization of biological processes (BPs) implicated by GO enrichment BP pathway analysis of DEGs in (**E)** anterior cingulate cortex, **(F)** amygdala, **(G)** nucleus accumbens and **(H)** caudate nucleus of *BAD1^+/−^* vs. WT pigs. BPs are grouped into gene networks based on functional similarity and shared genes, illustrating their interconnectivity. Node size reflects the number of genes in each BP. **I.** Reactome pathway enrichment analysis of DEGs across brain regions in *BAD1^+/−^* and WT pigs. **J.** Joint pathway analysis integrating transcriptomic and metabolomic datasets from each brain region. Differentially expressed genes and nominally altered metabolites (p < 0.05) were analyzed using the Joint Pathway Analysis module in MetaboAnalyst. Bubble size indicates pathway impact based on pathway topology analysis, and color denotes the average log(fold change) of pathway-associated features.

To assess whether these transcriptomic signatures were accompanied by changes in neurotransmitter levels or biochemical metabolic alterations, we next performed targeted neurotransmitter measurements from the same tissue biopsies. However, no genotype-dependent differences were observed in dopamine or serotonin levels across the investigated regions (**Figure S14A-B**). We then extended the analysis to untargeted metabolomics on the same brain tissues. Although no single metabolite reached FDR significance, several compounds were nominally altered (p < 0.05) in a region-specific manner (**Table S7-S10**). Metabolic pathway enrichment analyses identified alterations related to glutathione metabolism, amino acid metabolism, phospholipid and plasmalogen biosynthesis, nucleotide metabolism, and cellular redox regulation (**Figure S15A-C)**. Notably, glutathione-related metabolites, including oxidized glutathione species, showed consistent dysregulation across multiple brain regions (**Figure S16**).

To determine whether transcriptomic and metabolomic alterations converged on common biological processes, we performed integrated pathway analysis across molecular layers. This revealed remarkable agreement between independent omics datasets, identifying recurrent perturbations of mitochondrial energy metabolism, redox regulation, phospholipid biology, amino acid metabolism, and carbohydrate metabolism across multiple brain regions (**Figure 4J**). Particularly prominent were pathways related to glycolysis/gluconeogenesis, the tricarboxylic acid cycle, glutathione metabolism, and phospholipid metabolism. The convergence of transcriptomic and metabolomic alterations across multiple brain regions provides orthogonal evidence that BRD1 haploinsufficiency disrupts coordinated neuro-metabolic homeostasis, with mitochondrial function, redox regulation, and cellular metabolism emerging as core biological processes consistently affected by reduced BRD1 function.

### *BRD1* haploinsufficiency perturbs systemic metabolic homeostasis

To investigate whether metabolic alterations associated with *BAD1* haploinsufficiency extended beyond the brain, we profiled the blood metabolome longitudinally at 6, 9, and 14 months of age. Although no metabolites remained significant after correction for multiple testing, sucrose levels were nominally elevated in *BAD1*^+/−^ pigs compared to WT controls (**Table S11**). Supporting altered systemic carbohydrate metabolism, *BAD1*^+/−^ pigs additionally displayed a modest but consistent reduction in peripheral blood glucose levels at 14 months of age, measured independently using a clinical glucose meter (**Figure 5A**).

**Figure 5.**
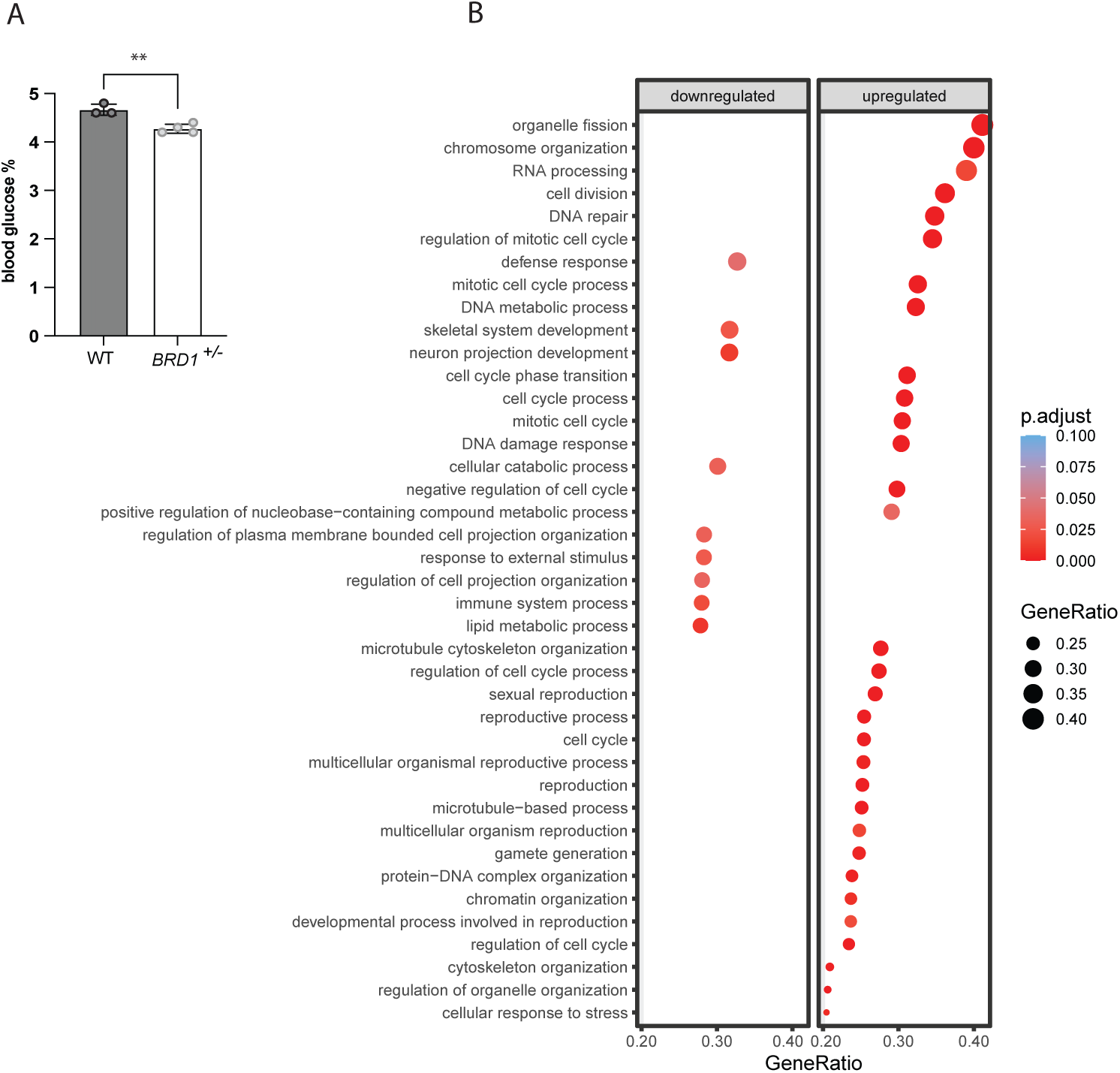
**A.** Peripheral blood glucose levels in *BAD1*^+/−^ and WT pigs at 14 months of age. Bars represent genotype mean±SD. **B.** Pathway enrichment analysis of DEGs in fibroblasts of *BAD1*^+/−^vs. WT pigs, sorted into downregulated vs. upregulated genes.

To further characterize systemic metabolic changes, we next analyzed the serum lipidome across the same developmental timepoints. Although no individual lipid species reached FDR significance, several phospholipid species were consistently altered across ages, including lipids corresponding to phosphatidylcholine and phosphatidylethanolamine species (**Table S12**).), which represent major structural components of cellular and mitochondrial membranes.

Finally, to determine whether systemic alterations were accompanied by peripheral cellular transcriptional changes, we profiled fibroblasts cultured from ear biopsies of newborn F1 pigs. Differential expression analysis identified 52 DEGs, including genes involved in cellular proliferation and metabolic regulation, such as *MI57* (**Table S13**). Pathway enrichment analysis further implicated cytoskeletal organization and metabolic regulatory processes (**Figure 5B**), supporting the notion that the molecular consequences of BRD1 haploinsufficiency extend beyond the central nervous system.

## Discussion

Understanding how psychiatric risk genes contribute to disease-relevant biology across multiple levels of organization remains a major challenge in psychiatric research. BRD1 has emerged as a compelling candidate in this context due to its repeated implication in psychiatric disorders and its central role in chromatin-mediated transcriptional regulation. To investigate the biological consequences of reduced BRD1 function in a translationally relevant system, we combined longitudinal neuroimaging with behavioral and multi-omics profiling in a genetically modified minipig model. BRD1 haploinsufficiency was associated with coordinated structural, molecular, and metabolic alterations across central and peripheral tissues.

While behavioral effects were subtle, longitudinal brain imaging revealed region-specific alterations in brain structure, and multi-omics profiling identified convergent molecular signatures consistent with altered mitochondrial and redox-related processes, phospholipid metabolism, and RNA processing. Together, these findings suggest that BRD1 influences interconnected neurodevelopmental and metabolic processes relevant to psychiatric disease biology.

While no gross developmental abnormalities were observed, *BAD1*^+/−^ pigs displayed a subtle age-dependent reduction in sucrose preference, suggesting altered motivational behavior. This finding is broadly consistent with previous observations in female *Brd1*^+/−^ rodent models, which exhibit affective and cognitive abnormalities, and supports the translational relevance of the porcine model. However, behavioral effects were modest compared to the more pronounced structural and molecular phenotypes observed across brain and peripheral tissues.

Neuroanatomically, *BAD1* haploinsufficiency was associated with region-specific differences in brain structure, particularly involving the cerebral cortex and caudate nucleus. Although total brain volume was not significantly affected, the observed pattern of regional volumetric differences suggests altered neurodevelopmental trajectories rather than generalized growth impairment. Structural abnormalities affecting cortical and striatal regions are among the most consistently reported neuroimaging findings across psychiatric and neurodevelopmental disorders, although the direction and timing of these changes vary considerably between conditions and developmental stages groups (63). For example, cortical thinning is frequently observed in schizophrenia, major depressive disorder, and bipolar disorder (64), whereas increased cortical thickness and surface area have been reported during early development in autism spectrum disorder (65). Similarly, enlarged caudate volume is a replicated finding in several neuroimaging studies of autism spectrum disorder (66–68). Notably, common genetic variation in *BAD1* has been associated with cerebral cortical surface area in large-scale human neuroimaging genetics studies (69), while *Brd1*^+/−^ mice exhibit structural brain abnormalities including reduced cortical and striatal volumes (69). Collectively, these findings support a role for BRD1 in regulating region-specific neurodevelopmental processes that contribute to structural brain variation and psychiatric disease-associated biology.

Consistent with its established role as a chromatin-associated transcriptional co-regulator, BRD1 haploinsufficiency produced substantial region-specific alterations in gene expression, with the most pronounced effects observed in the amygdala. Differentially expressed genes included regulators of excitatory and inhibitory neurotransmission, synaptic signaling, and hormone-responsive transcriptional programs. More broadly, enrichment analyses identified recurring signals related to oxidative phosphorylation, RNA processing, translation, and neural system function across multiple brain regions. Importantly, these transcriptomic signatures were independently supported by metabolomic profiling. Integrated analysis of transcriptomic and metabolomic datasets revealed striking convergence across molecular layers on pathways related to mitochondrial energy metabolism, glutathione metabolism, phospholipid biology, RNA processing, and translational regulation. Particularly notable was the repeated enrichment of selenoamino acid and glutathione metabolism. Seleno-proteins, including glutathione peroxidases and thioredoxin reductases, play central roles in antioxidant defense and maintenance of mitochondrial integrity and have been implicated in both neurodevelopment and psychiatric disease vulnerability (70).

The convergence of independent molecular modalities across multiple brain regions suggests that BRD1 haploinsufficiency disrupts coordinated neuro-metabolic homeostasis rather than isolated molecular pathways. This interpretation is further supported by previous experimental work demonstrating impaired mitochondrial respiration and altered bioenergetics following BRD1 deficiency in human neuronal models (19), suggesting that mitochondrial dysfunction may represent a conserved downstream consequence of reduced BRD1 function. Given the exceptionally high energetic demands of neurons, even relatively subtle disturbances in mitochondrial function and cellular energy metabolism may disproportionately affect neurodevelopment, synaptic function, and neuronal homeostasis. More broadly, converging evidence from genetic, transcriptomic, and postmortem studies has increasingly implicated mitochondrial dysfunction, oxidative stress, and altered cellular energy metabolism across schizophrenia, bipolar disorder, and major depressive disorder (40). Together, these findings support the emerging concept that psychiatric risk genes may contribute to disease biology through disruption of fundamental cellular processes required for normal neuronal development and function.

The effects of BRD1 haploinsufficiency were not confined to the central nervous system. Peripheral metabolomic and lipidomic profiling revealed signatures consistent with altered carbohydrate and lipid metabolism, including reduced circulating glucose levels and changes in phosphatidylcholine and phosphatidylethanolamine species. Although individual molecular changes were modest, the recurrence of related metabolic signatures across independent assays supports broader metabolic consequences of BRD1 disruption. Transcriptional alterations observed in cultured fibroblasts further suggest that BRD1-associated molecular dysregulation extends beyond neural tissues, consistent with its broad regulatory role across cell types.

The observation that BRD1-associated molecular alterations extend beyond the central nervous system may also be relevant in light of the growing recognition that psychiatric disorders are frequently accompanied by metabolic comorbidities, including insulin resistance, type 2 diabetes, obesity, and cardiovascular disease. Although lifestyle factors and psychotropic medication contribute substantially to this burden, accumulating evidence suggests that shared biological mechanisms also play a role. While the present study was not designed to model metabolic disease, the coordinated alterations observed across brain and peripheral tissues raise the possibility that BRD1 contributes to biological processes linking psychiatric vulnerability with systemic metabolic dysfunction.

Several limitations should, however, be considered when interpreting the present findings. First, the study was conducted in a relatively small cohort and included only female animals, limiting statistical power and preventing assessment of sex-dependent effects. Second, while transcriptomic signatures and pathway-level analyses showed consistent convergence across molecular layers, many individual metabolite and lipid measurements did not remain significant following correction for multiple testing and should therefore be interpreted cautiously. Nevertheless, the recurrence of related biological pathways across independent molecular modalities strengthens confidence in the broader biological themes identified.

In summary, *BAD1* haploinsufficiency was associated with coordinated neurodevelopmental and metabolic alterations across brain and peripheral tissues in a translational large-animal model. By integrating behavioral, neuroanatomical, transcriptomic, and metabolomic data, this study provides systems-level insight into how reduced BRD1 function disrupts interconnected neurodevelopmental and metabolic processes across multiple levels of biological organization. These findings support a role for BRD1 in coordinating molecular pathways involved in cellular homeostasis and illustrate how perturbation of a psychiatric risk gene can influence both brain function and systemic physiology.

## Acknowledgments

We extend our special thanks to Trine Werenberg Mikkelsen at Core Centre for Molecular Morphology, Section for Stereology and Microscopy, Aarhus University, for providing indispensable technical and practical assistance in immunostaining and tissue sectioning of minipig brains and to Lauren Van den Wijngaert for validating antibodies and optimizing sectioning protocol. We thank Ellegaard Göttingen Minipigs A/S for their support, and we further thank Martin Agergaard Fredsted, Mette Harder Bak, and Anne Mette Viernfeldt Toft for their dedicated care of the animals and for their invaluable assistance with animal handling, blood sampling, transport, and setup of behavioral testing. The project was supported by the A. P. Moeller Foundation, Helga og Peter Kornings Fond and Torben & Alice Frimodts Fond. The iPSYCH team was supported by grants from the Lundbeck Foundation (R102-A9118, R155-2014-1724, and R248-2017-2003).

## Author contributions

Conceptualization: PQ, ADB

Methodology: JGD, TF, BN, SHlC, SP, LZ, PRM, DPM, JEH, AM, JHC, IEH, GW, SFE, TEL, DG, JRN, FO, ME, AKOA, JJ, ADB, PQ

Investigation: JGD, PQ

Visualization: JGD, PRM, DPM, PQ, SFE

Funding acquisition: JGD, PQ, ADB

Project administration: JGD, PQ,

Supervision: PQ, ADB

Writing – original draft: JGD, PQ

Writing – review & editing: All authors

## References

1. Andreassen OA, Hindley GFLL, Frei O, Smeland OB. New insights from the last decade of research in psychiatric genetics: discoveries, challenges and clinical implications. World Psychiatry. 2023 Feb 14;22(1):4–24. doi:10.1002/wps.21034 PubMed PMID: 36640404.

2. Aberg K a, Liu Y, Bukszár J, McClay JL, Khachane AN, Andreassen O a, et al. A comprehensive family-based replication study of schizophrenia genes. Vol. 70. 2013 Feb;70(2). doi:10.1001/jamapsychiatry.2013.288

3. Nyegaard M, Severinsen JE, Als TD, Hedemand A, Straarup S, Nordentoft M, et al. Support of association between BRD1 and both schizophrenia and bipolar affective disorder. Am J Med Genet B Neuropsychiatr Genet. 2010 Mar;153B(2):582–91. doi:10.1002/ajmg.b.31023 PubMed PMID: 19693800.

4. Severinsen JE, Bjarkam CR, Kiaer-Larsen S, Olsen IM, Nielsen MM, Blechingberg J, et al. Evidence implicating BRD1 with brain development and susceptibility to both schizophrenia and bipolar affective disorder. Mol Psychiatry. 2006 Dec;11(12):1126–38. doi:10.1038/sj.mp.4001885 PubMed PMID: 16924267.

5. Jorgensen TH, Borglum AD, Mors O, Wang AG, Pinaud M, Flint TJ, et al. Search for Common Haplotypes on Chromosome 22q in Patients With Schizophrenia or Bipolar Disorder From the Faroe Islands. Am J Med Genet. 2002;114(2):245–52. doi:10.1002/ajmg.10191

6. Pardiñas AF, Holmans P, Pocklington AJ, Escott-Price V, Ripke S, Carrera N, et al. Common schizophrenia alleles are enriched in mutation-intolerant genes and in regions under strong background selection. Nat Genet. 2018 Mar 26;50(3):381–9. doi:10.1038/s41588-018-0059-2

7. Andreassen OA, Thompson WK, Dale AM. Boosting the power of schizophrenia genetics by leveraging new statistical tools. Schizophr Bull. 2014;40(1):13–7. doi:10.1093/schbul/sbt168 PubMed PMID: 24319118.

8. Bigdeli TB, Chatzinakos C, Bendl J, Barr PB, Venkatesh S, Gorman BR, et al. Biological Insights from Schizophrenia-associated Loci in Ancestral Populations. medRxiv. 2024 Aug 28;17:2024.08.27.24312631. doi:10.1101/2024.08.27.24312631

9. Purcell SM, Moran JL, Fromer M, Ruderfer D, Solovieff N, Roussos P, et al. A polygenic burden of rare disruptive mutations in schizophrenia. Nature. 2014;506(7487):185–90. doi:10.1038/nature12975 PubMed PMID: 24463508.

10. Iossifov I, O’Roak BJ, Sanders SJ, Ronemus M, Krumm N, Levy D, et al. The contribution of de novo coding mutations to autism spectrum disorder. Nature. 2014 Nov 29;515(7526):216–21. doi:10.1038/nature13908

11. Sanders SJ, Murtha MT, Gupta AR, Murdoch JD, Raubeson MJ, Willsey AJ, et al. De novo mutations revealed by whole-exome sequencing are strongly associated with autism. Nature. 2012 May 4;485(7397):237–41. doi:10.1038/nature10945

12. Xu LM, Li JR, Huang Y, Zhao M, Tang X, Wei L. AutismKB: an evidence-based knowledgebase of autism genetics. Nucleic Acids Res. 2012 Jan;40(D1):D1016–22. doi:10.1093/nar/gkr1145

13. Azidane S, Gallego X, Durham L, Cáceres M, Guney E, Pérez-Cano L. Identification of novel driver risk genes in CNV loci associated with neurodevelopmental disorders. Human Genetics and Genomics Advances. 2024 Jul;5(3):100316. doi:10.1016/j.xhgg.2024.100316

14. Doyon Y, Cayrou C, Ullah M, Landry AJ, Côté V, Selleck W, et al. ING tumor suppressor proteins are critical regulators of chromatin acetylation required for genome expression and perpetuation. Mol Cell. 2006 Jan;21(1):51–64. doi:10.1016/j.molcel.2005.12.007 PubMed PMID: 16387653.

15. Mishima Y, Miyagi S, Saraya A, Negishi M, Endoh M, Endo TA, et al. The Hbo1-Brd1/Brpf2 complex is responsible for global acetylation of H3K14 and required for fetal liver erythropoiesis. Blood. 2011 Sep;118(9):2443–53. doi:10.1182/blood-2011-01-331892 PubMed PMID: 21753189.

16. Fryland T, Christensen JH, Pallesen J, Mattheisen M, Palmfeldt J, Bak M, et al. Identification of the BRD1 interaction network and its impact on mental disorder risk. Genome Med. 2016 Dec;8(1):53. doi:10.1186/s13073-016-0308-x

17. McCullagh P, Chaplin T, Meerabux J, Grenzelias D, Lillington D, Poulsom R, et al. The cloning, mapping and expression of a novel gene, BRL, related to the AF10 leukaemia gene. Oncogene. 1999 Dec 9;18(52):7442–52. doi:10.1038/sj.onc.1203117 PubMed PMID: 10602503.

18. Cho HI, Kim MS, Jang YK. The BRPF2/BRD1-MOZ complex is involved in retinoic acid-induced differentiation of embryonic stem cells. Exp Cell Res. 2016 Aug;346(1):30–9. doi:10.1016/j.yexcr.2016.05.022

19. Paternoster V, Cömert C, Kirk LS, la Cour SH, Fryland T, Fernandez-Guerra P, et al. The psychiatric risk gene BRD1 modulates mitochondrial bioenergetics by transcriptional regulation. Transl Psychiatry. 2022 Dec 1;12(1). doi:10.1038/S41398-022-02053-2, PubMed PMID: 35941107.

20. Bjarkam CR, Corydon TJ, Olsen IML, Pallesen J, Nyegaard M, Fryland T, et al. Further immunohistochemical characterization of BRD1 a new susceptibility gene for schizophrenia and bipolar affective disorder. Brain Struct Funct. 2009 Dec;214(1):37–47. doi:10.1007/s00429-009-0219-3 PubMed PMID: 19763615.

21. Colantuoni C, Lipska BK, Ye T, Hyde TM, Tao R, Leek JT, et al. Temporal dynamics and genetic control of transcription in the human prefrontal cortex. Nature. 2011 Oct;478(7370):519–23. doi:10.1038/nature10524

22. Severinsen JE, Bjarkam CR, Kiaer-Larsen S, Olsen IM, Nielsen MM, Blechingberg J, et al. Evidence implicating BRD1 with brain development and susceptibility to both schizophrenia and bipolar affective disorder. Mol Psychiatry. 2006 Dec;11(12):1126–38. doi:10.1038/sj.mp.4001885 PubMed PMID: 16924267.

23. Fryland T, Christensen JH, Pallesen J, Mattheisen M, Palmfeldt J, Bak M, et al. Identification of the BRD1 interaction network and its impact on mental disorder risk. Genome Med. 2016 Dec;8(1):53. doi:10.1186/s13073-016-0308-x

24. Donskov J, Deans P, Høgfeldt J, Borglum A, Denham M, Brennand K, et al. BRD1 haploinsufficiency alters early neuronal programming and disrupts maturation in human induced glutamatergic neurons. 2026. doi:10.21203/rs.3.rs-8417267/v1

25. Qvist Per, Christensen JH, Vardya Irina, Rajkumar AP, Mørk Arne, Paternoster Veerle, et al. The schizophrenia associated BRD1 gene regulates behavior, neurotransmission, and expression of schizophrenia risk enriched gene sets in mice. Biol Psychiatry. 2016 Sep. doi:10.1016/j.biopsych.2016.08.037

26. Qvist P, Rajkumar AP, Redrobe JP, Nyegaard M, Christensen JH, Mors O, et al. Mice heterozygous for an inactivated allele of the schizophrenia associated Brd1 gene display selective cognitive deficits with translational relevance to schizophrenia. Neurobiol Learn Mem. 2017 May;141:44–52. doi:10.1016/j.nlm.2017.03.009 PubMed PMID: 28341151.

27. Rajkumar, Anto P. Qvist, Per. Lazarus, Ross. Nava, Nicoletta. Winther, Gudrun. Liebenberg, Nico. Paternoster, Veerle. Mørk, Arne. Fryland, Tue. Nyegaard, Mette. Nyengaard, Jens R. Wegener, Gregers. Mors, Ole. Christensen, Jane H. Børglum AD. Female Brd1+/− mice display reversible depressive phenotype, neurochemistry, neuronal morphology, and transcriptome that are consistent with major depressive disorder. In preparation.

28. Rajkumar, Anto P. Qvist, Per. Lazarus, Ross. Nava, Nicoletta. Winther, Gudrun. Liebenberg, Nico. Paternoster, Veerle. Mørk, Arne. Fryland, Tue. Nyegaard, Mette. Nyengaard, Jens R. Wegener, Gregers. Mors, Ole. Christensen, Jane H. Børglum AD. Female Brd1+/− mice display reversible depressive phenotype, neurochemistry, neuronal morphology, and transcriptome that are consistent with major depressive disorder. In preparation.

29. Qvist P, Eskildsen SF, Hansen B, Ringgaard S, Stødkilde H, Mors O, et al. The implication of the schizophrenia-associated BRD1 gene in brain morphology.

30. Paternoster V, Cömert C, Kirk LS, la Cour SH, Fryland T, Fernandez-Guerra P, et al. The psychiatric risk gene BRD1 modulates mitochondrial bioenergetics by transcriptional regulation. Transl Psychiatry. 2022 Aug 8;12(1):319. doi:10.1038/s41398-022-02053-2

31. Wong HHW, Chou CYC, Watt AJ, Sjöström PJ. Comparing mouse and human brains. Elife. 2023 Jul 10;12. doi:10.7554/ELIFE.90017 PubMed PMID: 37428552.

32. Chen K, Baxter T, Muir WM, Groenen MA, Schook LB. Genetic resources, genome mapping and evolutionary genomics of the pig (Sus scrofa). Int J Biol Sci. 2007;3(3):153–65. PubMed PMID: 17384734.

33. Hofman MA. Size and shape of the cerebral cortex in mammals. I. The cortical surface. Brain Behav Evol. 1985 Jan;27(1):28–40. PubMed PMID: 3836731.

34. Holm IE, Geneser FA. Histochemical demonstration of zinc in the hippocampal region of the domestic pig: I. Entorhinal area, parasubiculum, and presubiculum. J Comp Neurol. 1989 Sep;287(2):145–63. doi:10.1002/cne.902870202 PubMed PMID: 2477401.

35. Thibault KL, Margulies SS. Age-dependent material properties of the porcine cerebrum: effect on pediatric inertial head injury criteria. J Biomech. 1998 Dec;31(12):1119–26. PubMed PMID: 9882044.

36. Lind N, Moustgaard A, Jelsing J. The use of pigs in neuroscience: modeling brain disorders. Neuroscience & …. 2007.

37. Dobbing J, Sands J. Comparative aspects of the brain growth spurt. Early Hum Dev. 1979 Mar;3(1):79–83. PubMed PMID: 118862.

38. Akerstedt T, Levi L. Circadian rhythms in the secretion of cortisol, adrenaline and noradrenaline. Eur J Clin Invest. 1978 Apr;8(2):57–8. PubMed PMID: 417934.

39. Bollen P, Ellegaard L. The Göttingen Minipig in Pharmacology and Toxicology. Pharmacol Toxicol. 1997 May 25;80(s2):3–4. doi:10.1111/j.1600-0773.1997.tb01980.x

40. Jacobsen KRosenmay, Alstrup AKOlsen, Hansen AKornerup, Bollen PJA. The laboratory pig [Internet]. CRC Press; 2025 [cited 2025 Sep 2]. Available from: https://pure.au.dk/portal/da/publications/the-laboratory-pig-2

41. Alstrup AKO, Winterdahl M. Imaging Techniques in Large Animals. Scandinavian Journal of Laboratory Animal Science. 2009 Dec 1;36(1):55–66. doi:10.23675/SJLAS.V36I1.169

42. ZiFit [Internet]. Available from: http://zifit.partners.org/ZiFiT/

43. Zhang Lab General Cloning Protocol [Internet]. Available from: https://media.addgene.org/cms/filer_public/6d/d8/6dd83407-3b07-47db-8adb-4fada30bde8a/zhang-lab-general-cloning-protocol-target-sequencing_1.pdf

44. Schmidt M, Kragh PM, Li J, Du Y, Lin L, Liu Y, et al. Pregnancies and piglets from large white sow recipients after two transfer methods of cloned and transgenic embryos of different pig breeds. Theriogenology. 2010 Oct;74(7):1233–40. doi:10.1016/j.theriogenology.2010.05.026

45. Coupe P, Yger P, Prima S, Hellier P, Kervrann C, Barillot C. An optimized blockwise non-local means denoising filter for 3-D magnetic resonance images. IEEE Trans Med Imaging. 2008 Apr;27(4):425–41. doi:10.1109/TMI.2007.906087, PubMed PMID: 18390341.

46. Sled JG, Zijdenbos AP, Evans AC. A nonparametric method for automatic correction of intensity nonuniformity in mri data. IEEE Trans Med Imaging. 1998;17(1):87–97. doi:10.1109/42.668698, PubMed PMID: 9617910.

47. Fonov V, Evans AC, Botteron K, Almli CR, McKinstry RC, Collins DL. Unbiased average age-appropriate atlases for pediatric studies. Neuroimage. 2011 Jan 1;54(1):313–27. doi:10.1016/j.neuroimage.2010.07.033 PubMed PMID: 20656036.

48. Conrad MS, Sutton BP, Dilger RN, Johnson RW. An in vivo three-dimensional magnetic resonance imaging-based averaged brain collection of the neonatal piglet (Sus scrofa). PLoS One. 2014 Sep 25;9(9). doi:10.1371/JOURNAL.PONE.0107650, PubMed PMID: 25254955.

49. Kim D, Langmead B, Salzberg SL. HISAT: A fast spliced aligner with low memory requirements. Nat Methods. 2015 Mar 31;12(4):357–60. doi:10.1038/NMETH.3317;TECHMETA=38,47;SUBJMETA=114,1647,1949,2785,514,631,794;KWRD=GENOME+INFORMATICS,RNA+SEQUENCING,SOFTWARE PubMed PMID: 25751142.

50. Pertea M, Pertea GM, Antonescu CM, Chang TC, Mendell JT, Salzberg SL. StringTie enables improved reconstruction of a transcriptome from RNA-seq reads. Nat Biotechnol. 2015 Feb 18;33(3):290–5. doi:10.1038/NBT.3122;TECHMETA=45,91;SUBJMETA=2019,212,2302,61,631;KWRD=GENOME+ASSEMBLY+ALGORITHMS,TRANSCRIPTOMICS PubMed PMID: 25690850.

51. Yu G, Wang LG, Han Y, He QY. ClusterProfiler: An R package for comparing biological themes among gene clusters. OMICS. 2012 May 1;16(5):284–7. doi:10.1089/OMI.2011.0118/ASSET/IMAGES/LARGE/FIGURE1.JPEG PubMed PMID: 22455463.

52. Schmid R, Heuckeroth S, Korf A, Smirnov A, Myers O, Dyrlund TS, et al. Integrative analysis of multimodal mass spectrometry data in MZmine 3. Nat Biotechnol. 2023 Apr 1;41(4):447–9. doi:10.1038/S41587-023-01690-2;SUBJMETA=114,1314,320,61,631;KWRD=DATA+PROCESSING,METABOLOMICS PubMed PMID: 36859716.

53. Wang M, Carver JJ, Phelan V V., Sanchez LM, Garg N, Peng Y, et al. Sharing and community curation of mass spectrometry data with Global Natural Products Social Molecular Networking. Nat Biotechnol. 2016 Sep 8;34(8):828–37. doi:10.1038/NBT.3597;TECHMETA=101,58;SUBJMETA=11,114,2398,296,631,638,639;KWRD=COMPUTATIONAL+PLATFORMS+AND+ENVIRONMENTS,MASS+SPECTROMETRY PubMed PMID: 27504778.

54. Nothias LF, Petras D, Schmid R, Dührkop K, Rainer J, Sarvepalli A, et al. Feature-based molecular networking in the GNPS analysis environment. Nat Methods. 2020 Sep 1;17(9):905–8. doi:10.1038/S41592-020-0933-6;SUBJMETA=114,2398,320,61,631,92;KWRD=COMPUTATIONAL+PLATFORMS+AND+ENVIRONMENTS,METABOLOMICS PubMed PMID: 32839597.

55. Dührkop K, Fleischauer M, Ludwig M, Aksenov AA, Melnik A V., Meusel M, et al. SIRIUS 4: a rapid tool for turning tandem mass spectra into metabolite structure information. Nat Methods. 2019 Apr 1;16(4):299–302. doi:10.1038/S41592-019-0344-8;SUBJMETA=114,1647,296,320,45,631,794;KWRD=MASS+SPECTROMETRY,METABOLOMICS,SOFTWARE PubMed PMID: 30886413.

56. Dührkop K, Nothias LF, Fleischauer M, Reher R, Ludwig M, Hoffmann MA, et al. Systematic classification of unknown metabolites using high-resolution fragmentation mass spectra. Nat Biotechnol. 2021 Apr 1;39(4):462–71. doi:10.1038/S41587-020-0740-8;SUBJMETA=114,1305,320,631,92;KWRD=MACHINE+LEARNING,METABOLOMICS PubMed PMID: 33230292.

57. Schulz M, Zieglowski L, Kopaczka M, Tolba RH. The Open Field Test as a Tool for Behaviour Analysis in Pigs: Recommendations for Set-Up Standardization – A Systematic Review. European Surgical Research. 2023 Mar 23;64(1):7–26. doi:10.1159/000525680

58. GWAS Catalog [Internet]. [cited 2025 May 2]. Available from: https://www.ebi.ac.uk/gwas/publications/35396580

59. Bristow GC, Bostrom JA, Haroutunian V, Sodhi MS. Sex differences in GABAergic gene expression occur in the anterior cingulate cortex in schizophrenia. Schizophr Res. 2015 Sep 1;167(1–3):57–63. doi:10.1016/J.SCHRES.2015.01.025 PubMed PMID: 25660468.

60. GWAS Catalog [Internet]. [cited 2025 May 2]. Available from: https://www.ebi.ac.uk/gwas/publications/30718901

61. GWAS Catalog [Internet]. [cited 2025 May 2]. Available from: https://www.ebi.ac.uk/gwas/publications/26198764

62. GWAS Catalog [Internet]. [cited 2025 May 2]. Available from: https://www.ebi.ac.uk/gwas/publications/34045744

63. Thompson PM, Jahanshad N, Ching CRK, Salminen LE, Thomopoulos SI, Bright J, et al. ENIGMA and global neuroscience: A decade of large-scale studies of the brain in health and disease across more than 40 countries. Transl Psychiatry. 2020;10(1):100. doi:10.1038/s41398-020-0705-1

64. Schmaal L, Hibar DP, Sämann PG, Hall GB, Baune BT, Jahanshad N, et al. Cortical abnormalities in adults and adolescents with major depression based on brain scans from 20 cohorts worldwide in the ENIGMA Major Depressive Disorder Working Group. Mol Psychiatry. 2017 Jun 1;22(6):900–9. doi:10.1038/MP.2016.60;TECHMETA=57,59;SUBJMETA=1414,476,692,699;KWRD=DEPRESSION PubMed PMID: 27137745.

65. Van Rooij D, Anagnostou E, Arango C, Auzias G, Behrmann M, Busatto GF, et al. Cortical and subcortical brain morphometry differences between patients with autism spectrum disorder and healthy individuals across the lifespan: Results from the ENIGMA ASD working group. American Journal of Psychiatry. 2018 Apr 1;175(4):359–69. doi:10.1176/APPI.AJP.2017.17010100, PubMed PMID: 29145754.

66. Langen M, Schnack HG, Nederveen H, Bos D, Lahuis BE, de Jonge M V., et al. Changes in the Developmental Trajectories of Striatum in Autism. Biol Psychiatry. 2009 Aug 15;66(4):327–33. doi:10.1016/j.biopsych.2009.03.017 PubMed PMID: 19423078.

67. Langen M, Bos D, Noordermeer SDS, Nederveen H, Van Engeland H, Durston S. Changes in the Development of Striatum Are Involved in Repetitive Behavior in Autism. Biol Psychiatry. 2014 Sep 1;76(5):405–11. doi:10.1016/J.BIOPSYCH.2013.08.013 PubMed PMID: 24090791.

68. Lange N, Travers BG, Bigler ED, Prigge MBD, Froehlich AL, Nielsen JA, et al. Longitudinal Volumetric Brain Changes in Autism Spectrum Disorder Ages 6–35 Years. Autism Research. 2015 Feb 1;8(1):82–93. doi:10.1002/AUR.1427 PubMed PMID: 25381736.

69. Shadrin AA, Kaufmann T, van der Meer D, Palmer CE, Makowski C, Loughnan R, et al. Vertex-wise multivariate genome-wide association study identifies 780 unique genetic loci associated with cortical morphology. Neuroimage. 2021 Dec 1;244:118603. doi:10.1016/j.neuroimage.2021.118603 PubMed PMID: 34560273.

70. Bjørklund G, Doşa MD, Maes M, Dadar M, Frye RE, Peana M, et al. The impact of glutathione metabolism in autism spectrum disorder. Pharmacol Res. 2021 Apr 1;166:105437. doi:10.1016/J.PHRS.2021.105437 PubMed PMID: 33493659.

