## Supplemental Figures for "BRD1 haploinsufficiency disrupts neurodevelopmental and metabolic homeostasis in a translational minipig model"

**Supplementary figures**

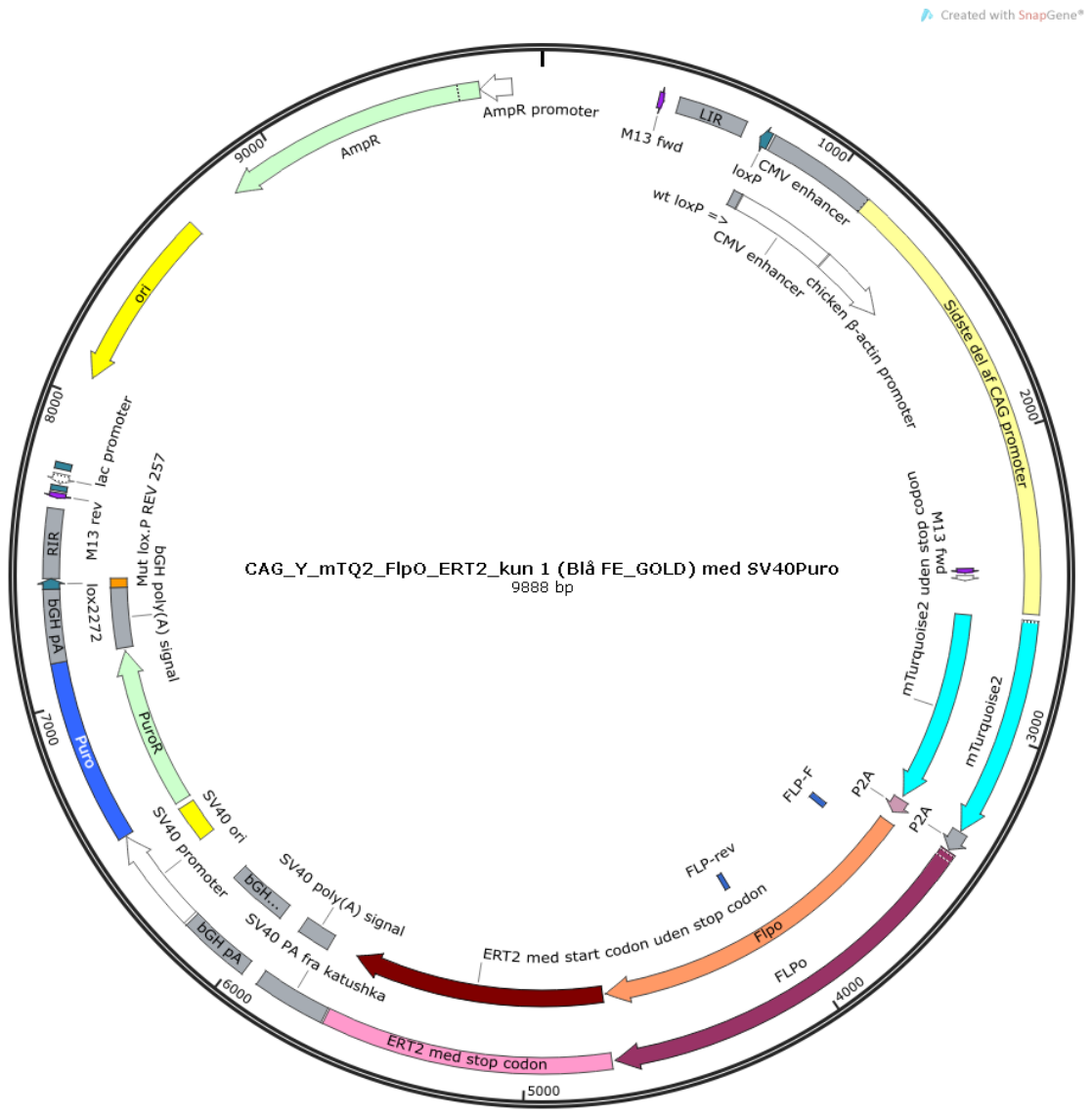

**Figure S1.** Blue-puro is the plasmid used with the fibroblast cells with a single transgene cassette containing the enhanced green fluorescent protein (eGFP) and is flanked with two heterozygous loxP sites. The fibroblasts were transfected with a plasmid encoding the Cre recombinase, and the plasmid shown here (Blue-puro) containing the puromycin resistant gene and it is flanked by the same heterozygous loxP sites.

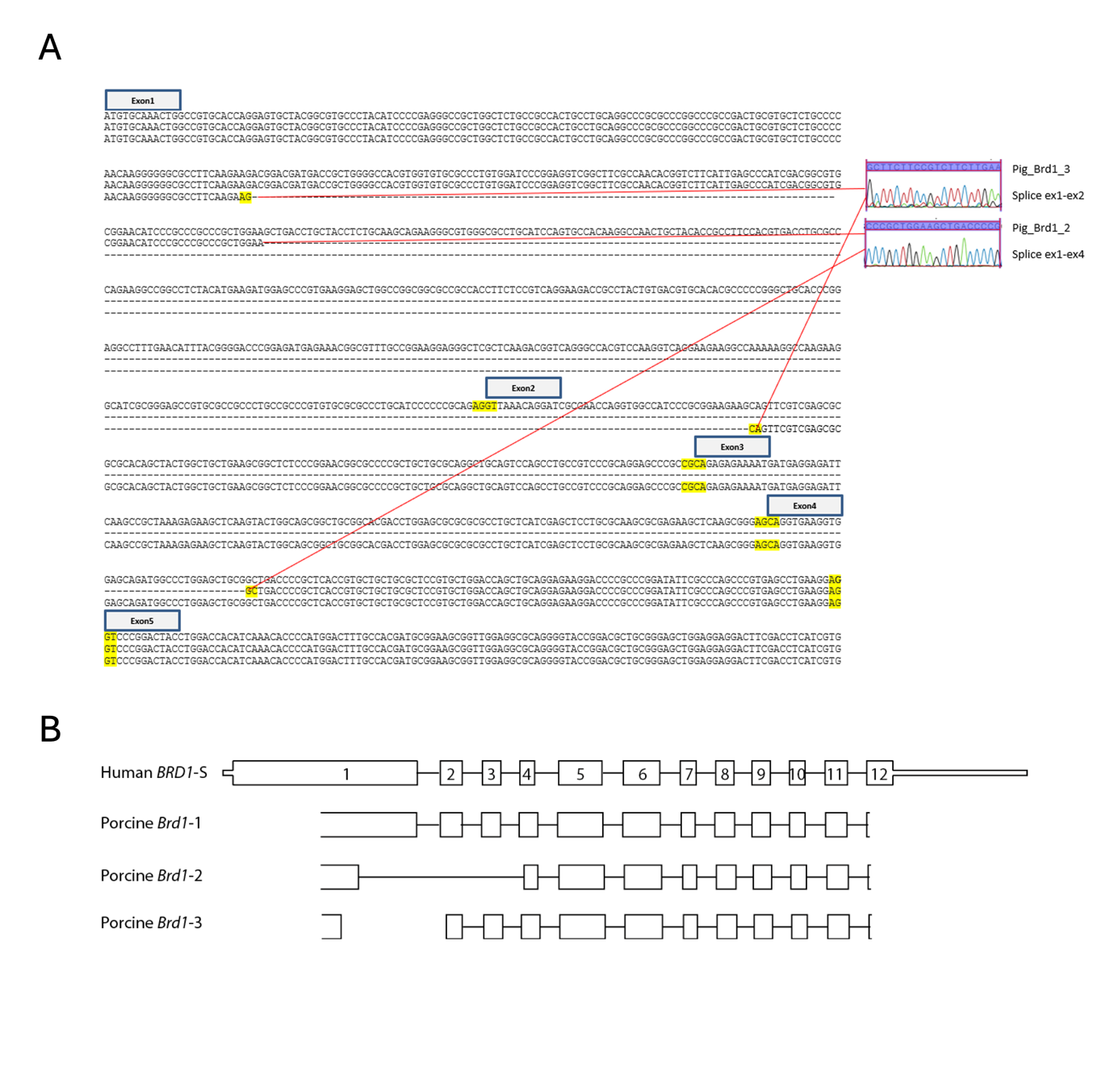

**Figure S2. A.** Exon-exon junctions in the coding sequence of *BRD1* in a Göttingen minipig. **B.** Schematic of the human short *BRD1* variant (BRD1-S) and porcine *BRD1* variants.

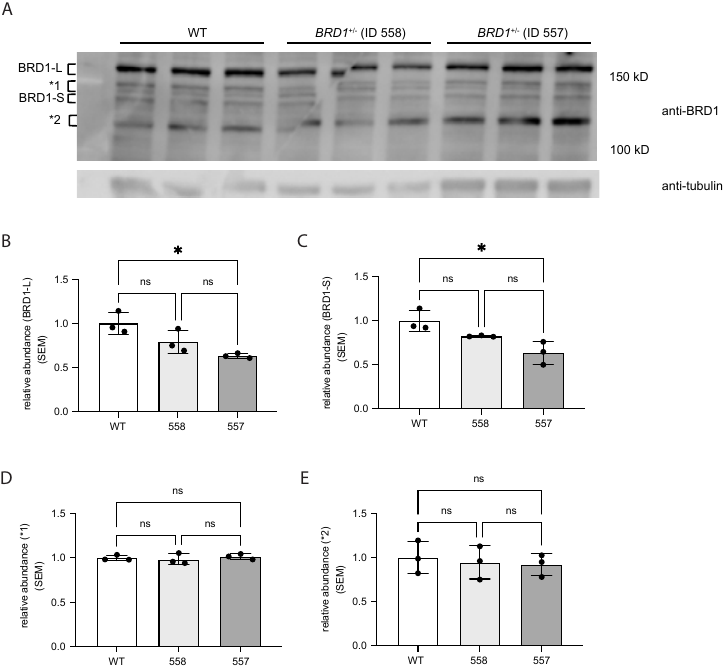

**Figure S3. BRD1 protein expression in minipig fibroblasts. A.** Immunoblotting was performed using custom-ordered affinity-purified chicken antibodies against BRD1 and secondary antibody (Horseradish Peroxidase Goat Anti-Chicken IgY H-1004). Tubulin was used as a loading control to normalize protein levels. **B–E.** Quantification of BRD1 protein. Plots show relative abundance in one WT and two *BRD1^+/-^* minipigs of isoform BRD1-L (B) and isoform BRD1-S (C). (D) and (E) show quantified unspecific binding. Bars represent mean±SD.

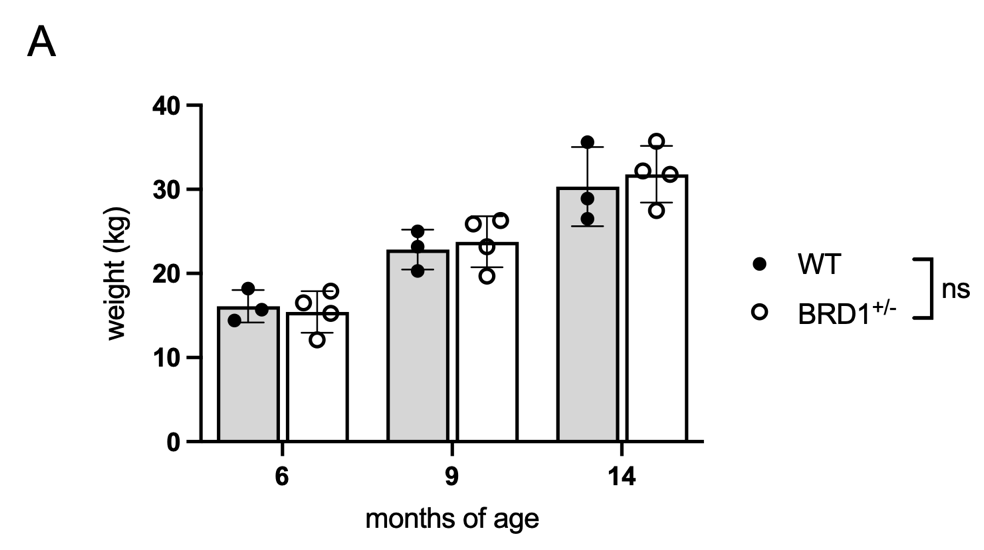

**Figure S4. Growth curves for WT and *BRD1^+/-^* minipigs throughout development.** Bars represent mean±SD.

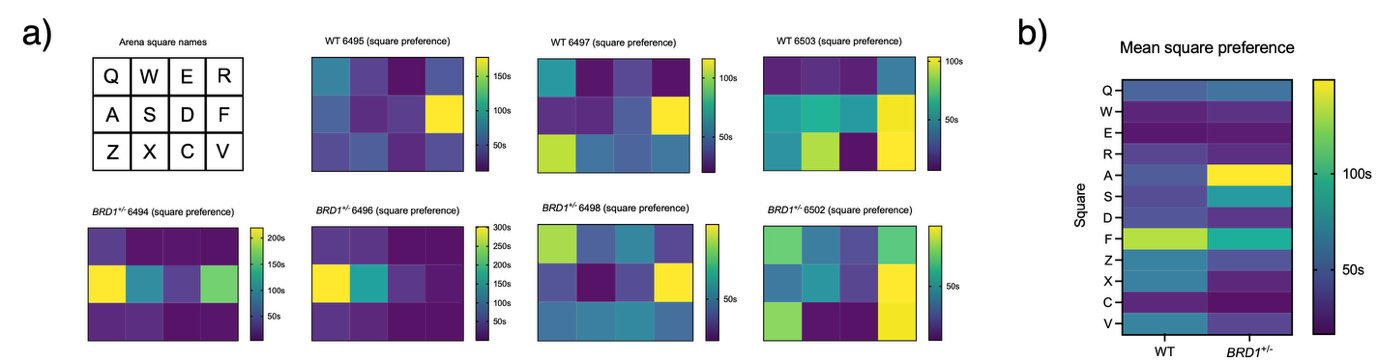

**Figure S5. OF novelty.** Arena location preference of WT (top) and *BRD1^+/-^* (bottom) minipigs. **A.** Illustration of arena, and the total time spent in each square. **B.** Group-mean of total time spent in each square.

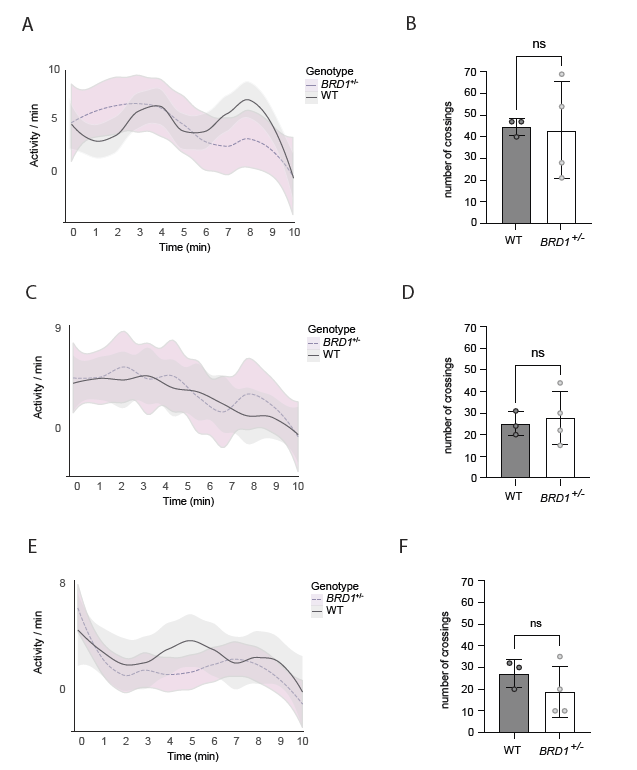

**Figure S6. Behavioral assessment of *BRD1^+/-^* and WT minipigs. A–F.** Activity of minipigs in open field (OF) test over time (A, C, E) or in total (B, D, F), measured by the number of crossings between squares. Graphs and bars represent genotype mean±SD. A–B: OF novelty, C–D: OF novelty+object, E–F: OF baseline.

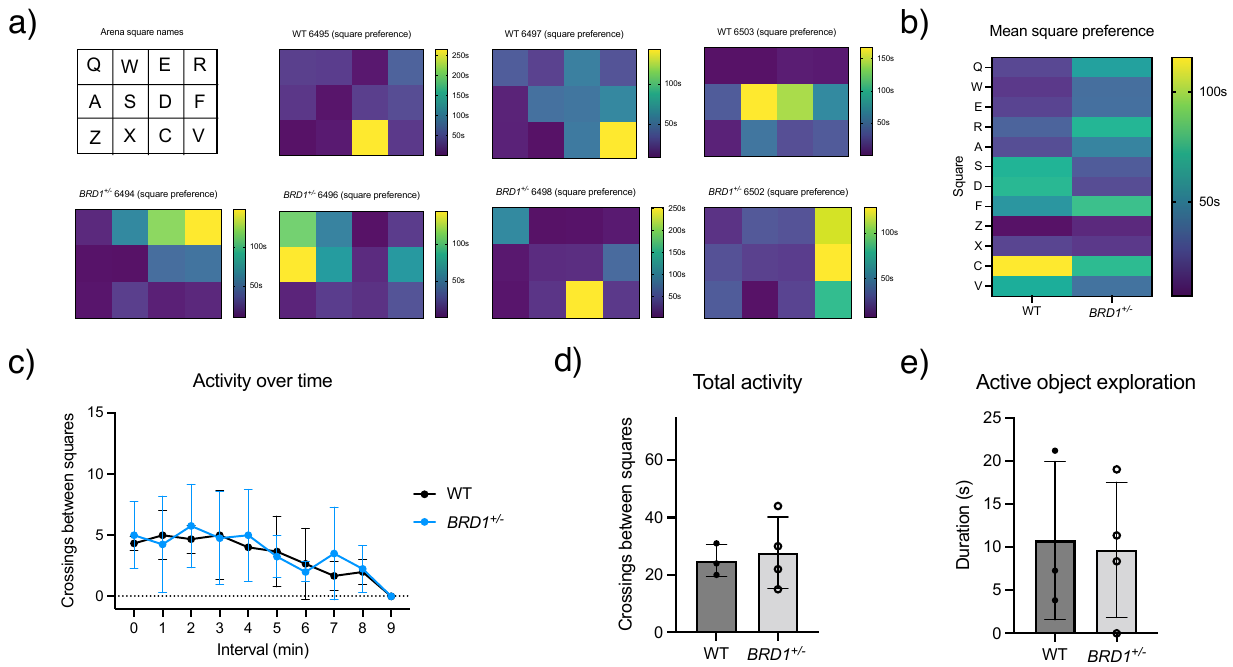

**Figure S7.** **OF novelty + object.** Arena location preference of WT (top) and *BRD1^+/-^* (bottom) minipigs. **A.** Illustration of arena, and the total time spent in each square. **B.** Group-mean of total time spent in each square.

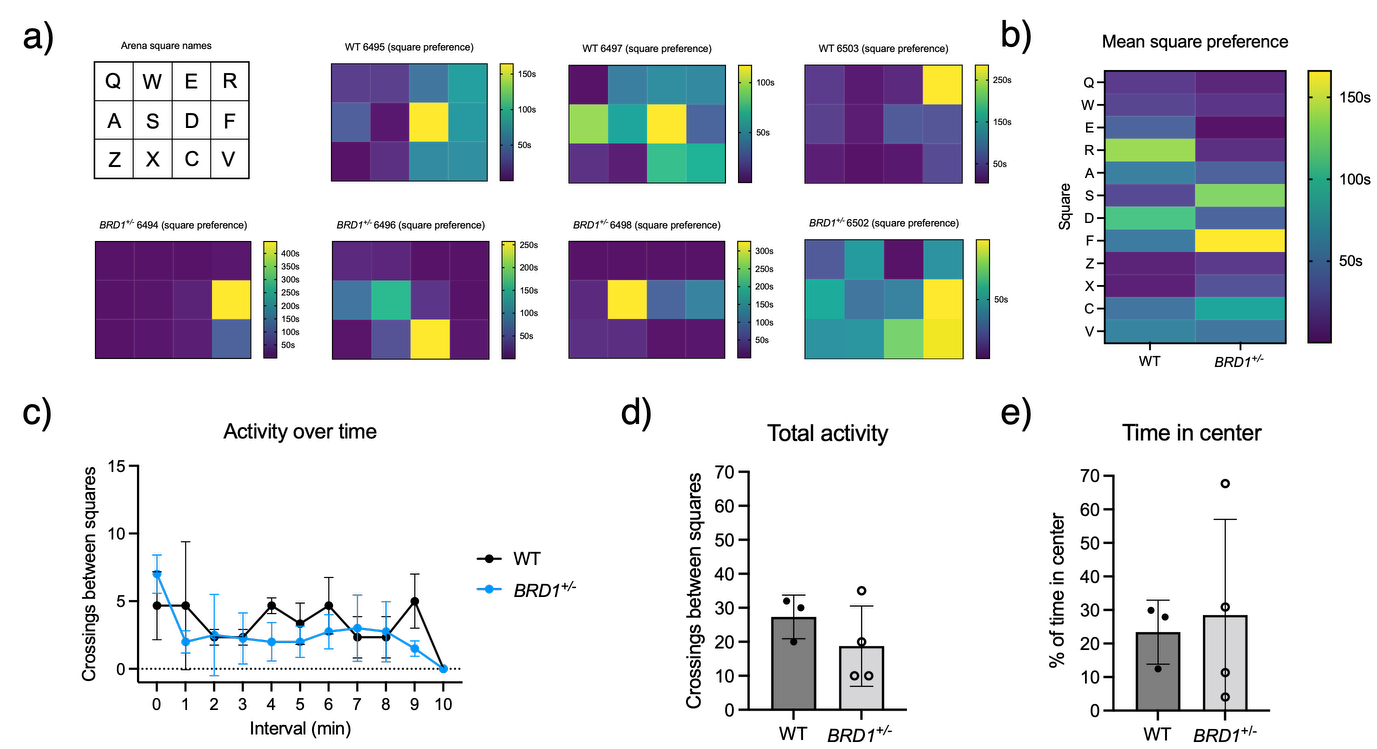

**Figure S8. OF baseline.** Arena location preference of WT (top) and *BRD1^+/-^* (bottom) minipigs. **A.** Illustration of arena, and the total time spent in each square. **B.** Group-mean of total time spent in each square.

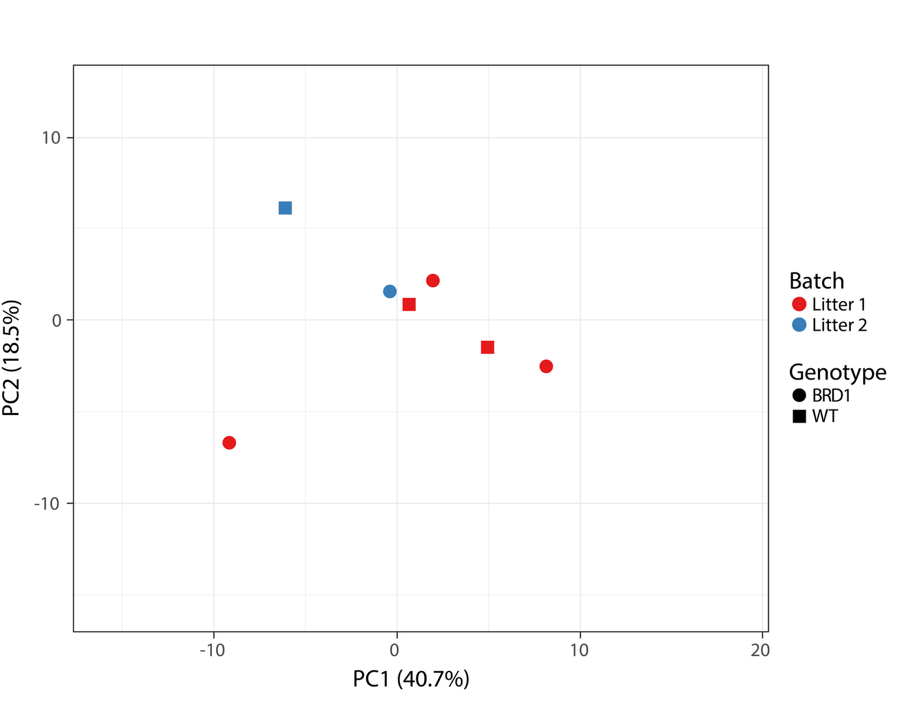

**Figure S9.** PCA plot of minipigs from litter 1 and litter 2 based on volumetric data from 20 brain regions.

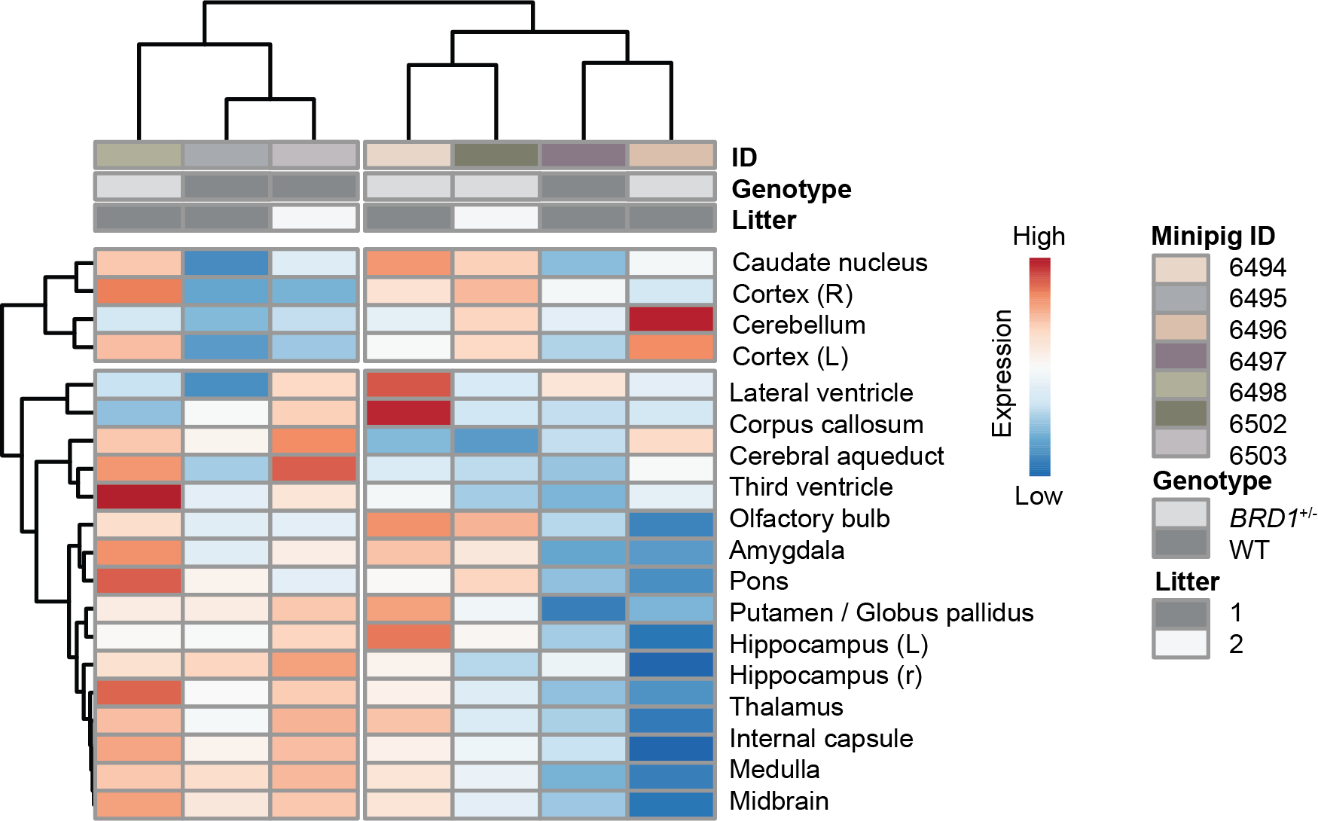

**Figure S10.** Heatmap showing hierarchical clustering of normalized and litter-corrected regional brain volumes at 14 months of age.

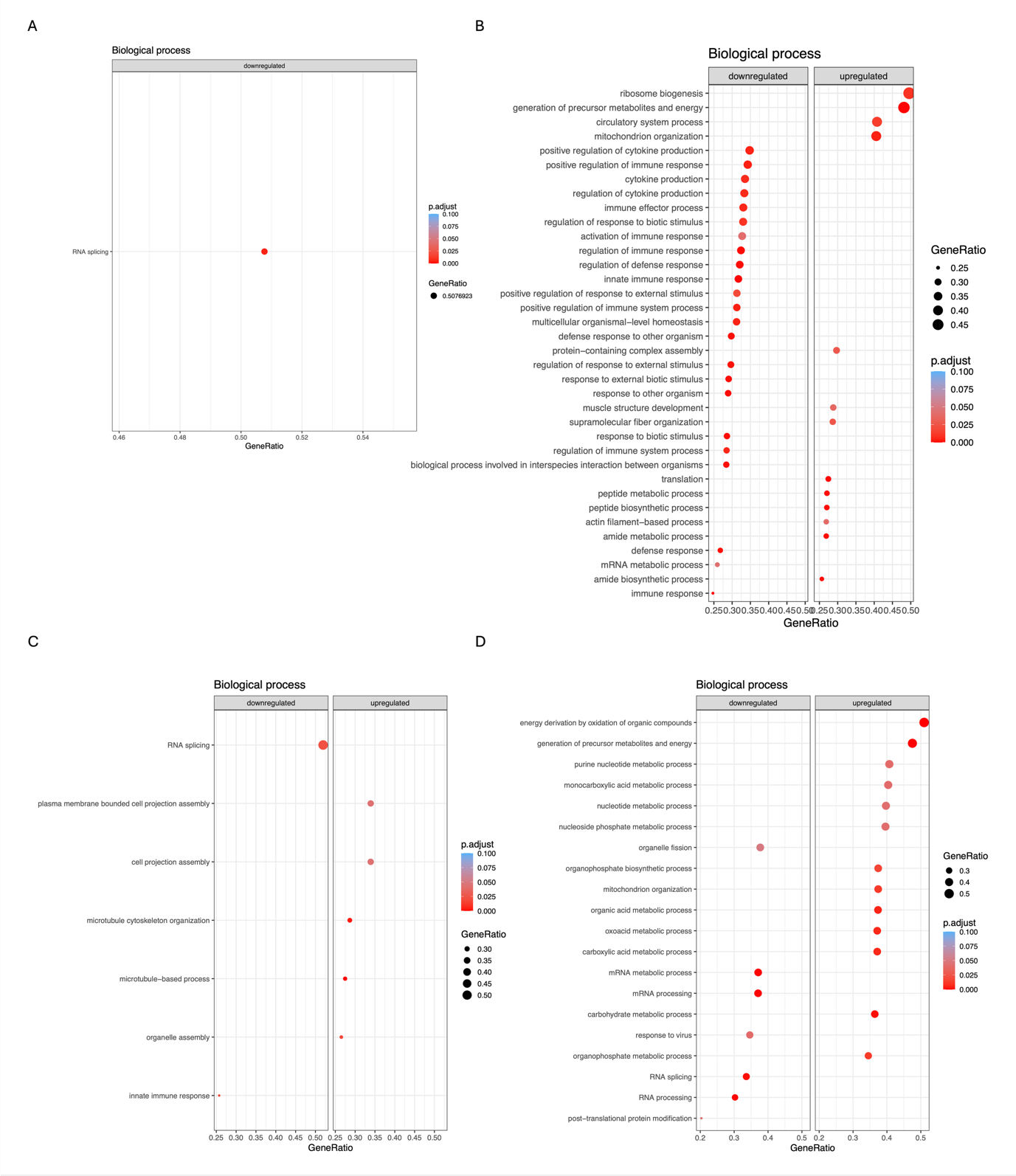

**Figure S11. GO biological process enrichment analysis of DEGs in brain regions. A.** AMG. **B.** aCC. **C.** NAc. **D.** NC.

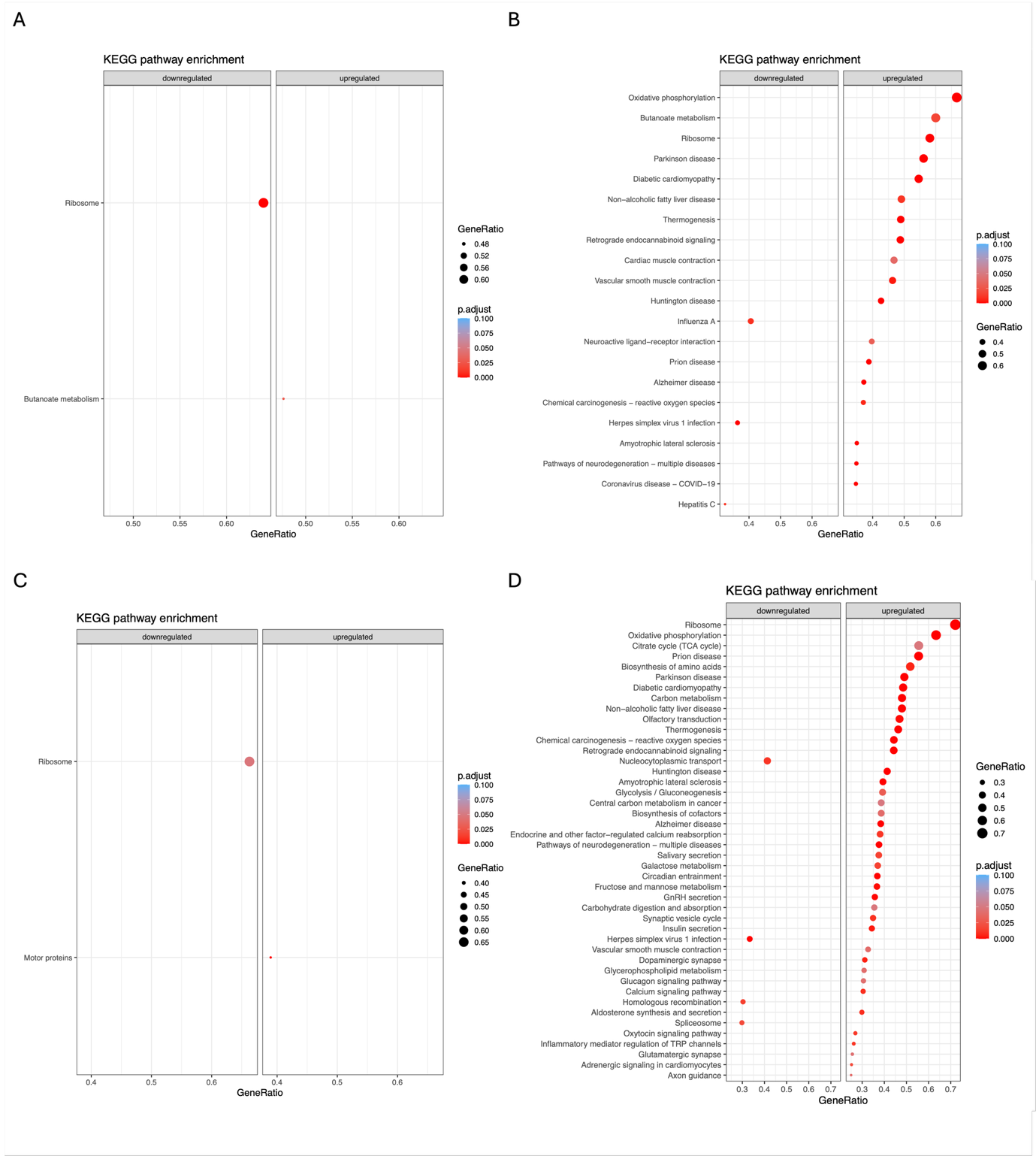

**Figure S12. KEGG pathway enrichment analysis of DEGs in brain regions. A.** AMG. **B.** aCC. **C.** NAc. **D.** NC.

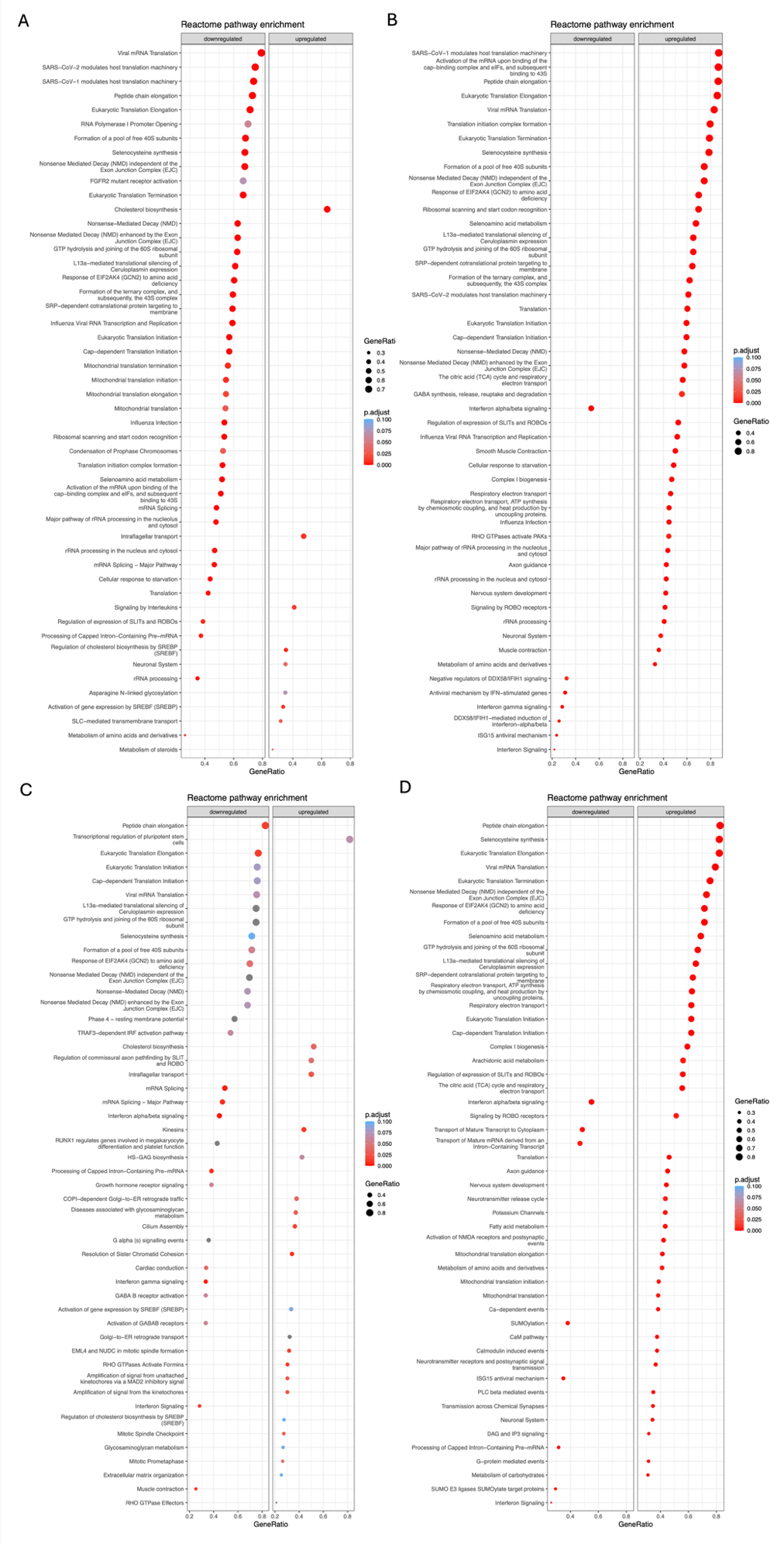

**Figure S13. Reactome pathway enrichment analysis of DEGs in brain regions. A.** AMG. **B.** aCC. **C.** NAc. **D.** NC.

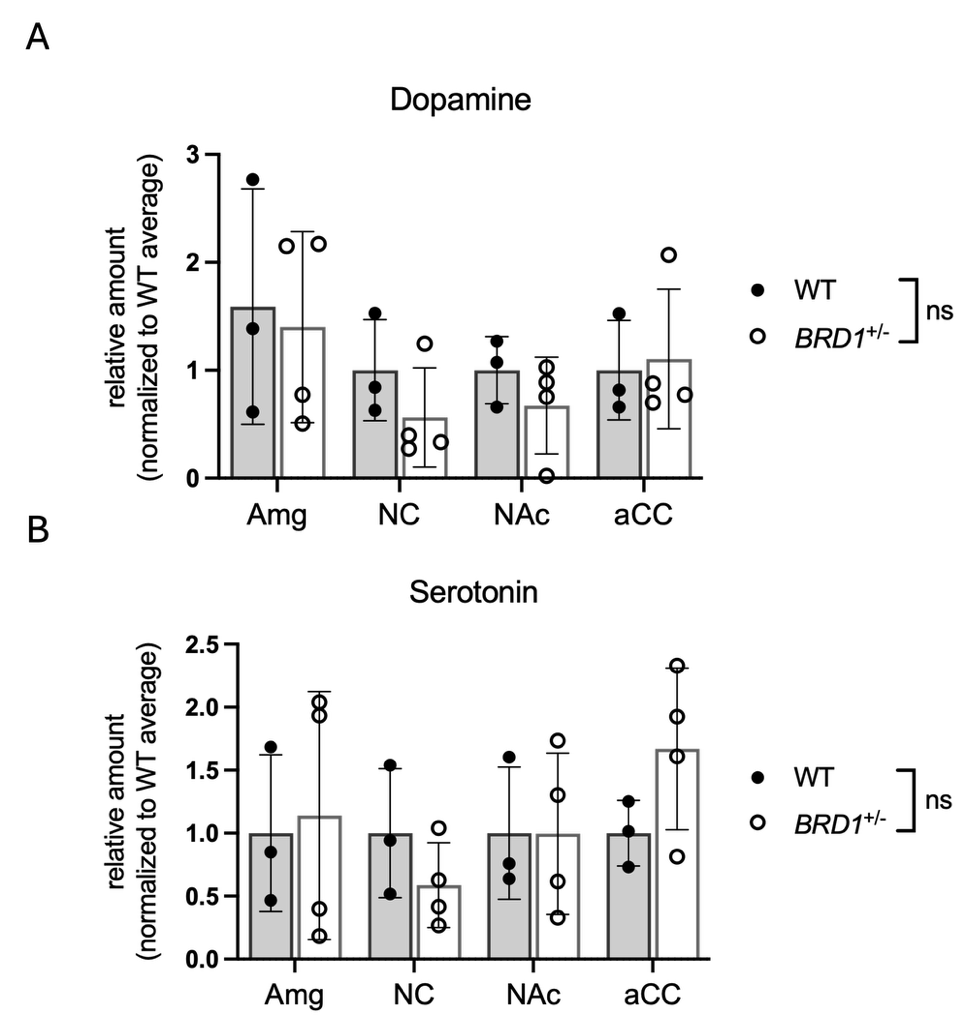

**Figure S14. Neurotransmitter levels in WT and *BRD1^+/-^* minipigs brains**. Bars represent mean±SD. **A.** Normalized dopamine levels. **B.** Normalized serotonin levels.

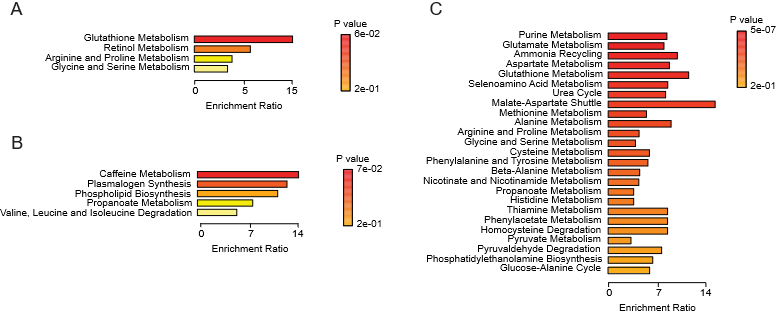

**Figure S15. Pathway enrichment analyses of altered metabolite levels in *BRD1^+/-^* minipigs.** **A.** aCC. **B.** AMG. **C.** NAc.

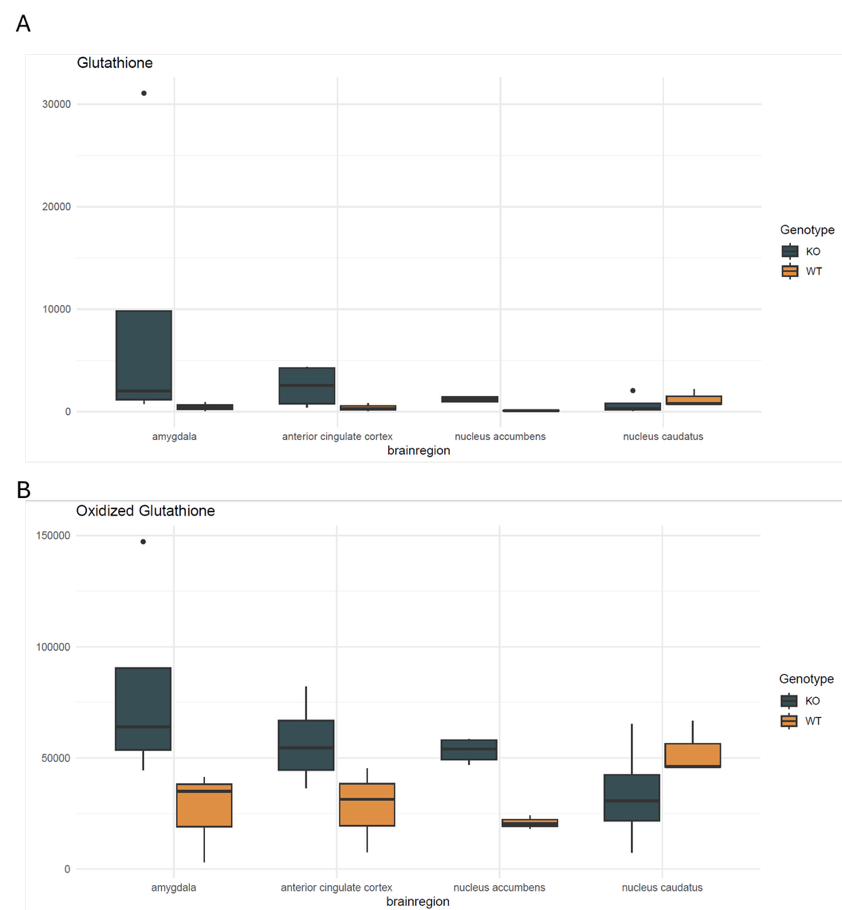

**Figure S16.** Levels of glutathione **(A)** and oxidized glutathione **(B)** in AMG, aCC, NAc and NC in WT and *BRD1^+/-^* minipigs.
